# Life without heterotrimeric kinesins: trypanosomatids use a combination of homodimeric kinesin-2 motors to drive intraflagellar transport

**DOI:** 10.64898/2026.05.12.724483

**Authors:** Aline Araujo Alves, Augustine Cleetus, Cécile Fort, Kristína Záhonová, Daniel Abbühl, Christine Girard-Blanc, Thierry Blisnick, Serge Bonnefoy, Nadège Cayet, Ziyin Wang, Jack D. Sunter, Vyacheslav Yurchenko, Richard J. Wheeler, Zeynep Ökten, Philippe Bastin

## Abstract

Heterotrimeric kinesin 2 is the canonical motor protein for anterograde intraflagellar transport (IFT), driving movement of protein complexes towards the tip of cilia and flagella. Here, we show that all members of the Euglenozoa group lack genes for heterotrimeric kinesins and instead possess a variable number of genes for two homodimeric kinesins termed KIN2A and KIN2B. When expressed *in vitro*, both *Trypanosoma brucei* kinesins form homodimers and move processively along brain microtubules, KIN2A being faster than KIN2B. Studies in *T. brucei* and *Leishmania mexicana* show anterograde and retrograde IFT of both kinesins, with KIN2A travelling throughout the whole length of the flagellum, while KIN2B is concentrated at its base. In the proximal portion of the flagellum, most KIN2B molecules travel without IFT proteins, except for a few particles that are associated with IFT proteins and reach the tip. Surprisingly, the absence of KIN2A has mild effects on IFT and flagellum assembly, whereas KIN2B is essential for both. Investigation of trypanosome flagella deprived of KIN2B revealed that IFT proteins do not access these flagella but that KIN2A can still circulate. We propose a division-of-labour model where KIN2B would be responsible for the import of IFT proteins while KIN2A would ensure most of the anterograde transport.

## Introduction

In eukaryotes, cilia and flagella are microtubule-based cell protrusions involved in various functions such as motility, sensing or morphogenesis [1]. They are separated from the rest of the cell body by the transition zone that acts as a filter to control protein entry and exit [2]. Cilia and flagella are constructed by a dynamic process called intraflagellar transport (IFT), where molecular motors drive IFT trains, large polymers made of two protein complexes (IFT-A and IFT-B), which shuttle precursors to the tip of the organelle where they are incorporated [3–6]. IFT is essential for cilia construction, and the 22 IFT proteins show remarkable conservation across eukaryotes [7].

Motors of the kinesin-2 family are responsible for anterograde transport, i.e. from the flagellum base to the tip. They associate with the IFT trains near the flagellum base and drag them through the transition zone, “walking” on the ciliary/flagellum axoneme microtubules [8, 9]. Kinesin-2 emerged together with cilia in evolution and consists of an amino-terminal motor domain followed by coiled-coil domains where cargoes and partners can dock [10]. The typical configuration is heterotrimeric kinesin-2, which is composed of two distinct motor subunits (termed KIF3A and KIF3B or KIF3A and KIF3C) and a kinesin-associated protein (KAP) that contains multiple armadillo repeats and a tyrosine-rich acidic carboxy-terminal tail but neither a microtubule-interacting motif nor a motor domain [10]. The heterotrimeric kinesin was first described in sea urchin, where it navigates towards the plus end of microtubules [11]. Seminal work in the green alga *Chlamydomonas reinhardtii* discovered that the *FLA10* gene encodes one of the two motor subunits and demonstrated kinesin-2 close association to IFT particles [12]. Inhibition of FLA10 resulted in arrest of IFT, accompanied by a block in new flagellum formation and a resorption of existing flagella [13], a finding reproduced in other organisms [14–16].

A second member of the kinesin-2 family was subsequently discovered in animals and termed OSM-3 or KIF17 [17, 18]. It is made of a different motor subunit that functions as a homodimer and does not need an accessory protein to ensure dimerization [19]. OSM-3 was studied extensively in the nematode *Caenorhabditis elegans*, where most sensory cilia are composed of three portions: a transition zone, a middle segment made of nine doublet microtubules and a distal segment composed of nine singlet microtubules. Absence of the homodimeric OSM-3 results in the formation of shorter cilia deprived of the distal segment while depletion of the heterotrimeric kinesin had no visible effect on cilium length [20]. Further studies demonstrated that the heterotrimeric kinesin ensures IFT train progression in the transition zone before being progressively substituted by OSM-3 during transport in the middle segment [8]. At the junction with the distal segment, the heterotrimeric kinesin is released from the trains and recycled to the base by retrograde transport [21]. Because OSM-3 is a faster motor, the speed of anterograde transport increases along ciliary length to reach its highest rate in the distal segment [21]. KIF17, the mammalian homologue of OSM-3, localises to cilia in neurons (especially photoreceptors) but is not required for their construction [22, 23]. However, it is involved in signalling and development in mouse [24], as well as in photoreceptors and olfactory neurons in zebrafish [25, 26].

The heterotrimeric kinesin is conserved in most eukaryotes assembling cilia and flagella using IFT. However, kinetoplastids, which belong to the Euglenozoa group [27, 28], lack KAP homolog [29, 30]. Instead, two genes encode very distinct kinesin-2 proteins called KIN2A and KIN2B that are predicted to be homodimeric [29]. Initial investigations were performed in *Trypanosoma brucei*, a group of parasites shuttling between tsetse flies and mammals and responsible for various human and animal diseases. Trypanosomes are also excellent model organisms to study the assembly of cilia and flagella [31] [32]. In the procyclic stage of the parasite (which develops in the midgut of the tsetse fly but can be cultivated and manipulated axenically), individual RNAi silencing of KIN2A or KIN2B did not lead to a visible IFT or flagellum phenotype, but a combined knockdown reduced both the rate and the speed of IFT trains, resulting in the assembly of shorter flagella [33]. In the trypanosome bloodstream stage (which develops in mammals and can also be manipulated), knockdown of KIN2A or KIN2B individually reduced flagellum length by 4-5 µm, and an additional reduction was observed with combined silencing [30]. These results suggest that the two kinesins may be redundant. Nevertheless, silencing of KIN2A alone, but not KIN2B, severely impacted the growth rate of bloodstream trypanosomes and led to spectacular cytokinesis defects [30], hinting at potentially some distinct functions. However, knockdown efficiency was less potent for KIN2B (∼30% residual mRNA vs ∼10% for KIN2A), hindering firm conclusions. It should also be reminded that bloodstream trypanosomes are more sensitive to flagellum perturbation compared to procyclic cells [34, 35].

The discovery of species assembling flagella and performing IFT without the canonical heterotrimeric kinesin-2 is intriguing and raises multiple questions: is the absence of KAP and the presence of two candidate homodimeric kinesins limited to kinetoplastids? Are KIN2A and KIN2B functioning as independent homodimers, or can they associate to form a previously unrecognised new type of heterodimeric kinesin motor? What is the capacity of these atypical kinesin motors for shuttling IFT proteins? And finally, do they function independently or do they cooperate?

Here, we performed extended *in silico* analyses and found that the absence of KAP and the presence of KIN2A and KIN2B are only found in all members of the phylum Euglenozoa, whereas other species retain KAP and distinct kinesin-2 genes. KIN2A and KIN2B of *T. brucei* form independent homodimers capable of moving along mammalian brain microtubules assembled *in vitro*. Endogenous tagging revealed distinct localisations in the flagellum: KIN2A moves along its entire length, while most KIN2B is restricted to the proximal region and only occasionally co-localises with IFT. Intriguingly, the absence of KIN2A has a mild effect on flagellum length despite disrupting IFT and redistributing KIN2B along the flagellum. Conversely, KIN2B deletion inhibits flagellum assembly. We propose a division-of-labour model in which KIN2B promotes the entry of IFT proteins or trains into the flagellum and KIN2A ensures most of their trafficking along the axoneme.

## Results

### KIN2A and KIN2B are restricted to the Euglenozoa group

The kinesins KIN2A and KIN2B of kinetoplastids are among the most divergent kinesin- 2 members identified to date [30, 36]. The genomes of all kinetoplastids lack a gene for KAP [29, 30], suggesting that these species could rely exclusively on homodimeric or possibly atypical heterodimeric kinesin-2. To understand the distribution of the IFT-related proteins in eukaryotes, we searched the sequencing data of 198 species (**Table S1**) for the presence of kinesin-2 coding genes (**Fig. 1A, Fig. S1**), as well as KAP and two of the most conserved IFT proteins, IFT52 and IFT88 (**Fig. 1B**) ([7]. To understand the relationships among the kinesin-2 sequences, we performed a comprehensive phylogenetic analysis (**Fig. 1A, Fig. S1**). Consistent with previous work [36], KIF3A, KIF3B, KIF3C, and KIF17 were specific for Holozoa (metazoans and their single-celled relatives). KIN2A and KIN2B were identified not only in kinetoplastids, but also in other members of the Euglenozoa phylum. However, while kinetoplastids contain single copies of each gene, several paralogous copies are present in other organisms. For example, the euglenids *Euglena gracilis* and *E. longa* possess three genes for KIN2A and one gene for KIN2B. The branching pattern indicates that KIN2A was multiplied in the common ancestor of the two euglenids. The diplonemid *Paradiplonema papillatum* (renamed from *Diplonema papillatum* [37]) has two copies of each. Notably, in the performed phylogenetic analysis, a sequence from the rhizarian alga *Bigelowiella natans* formed a highly supported (97 ultrafast bootstrap [UFB]) sister lineage to the euglenozoan monophyletic clade comprising all KIN2A and KIN2B sequences (100 UFB) (**Fig. 1A**). However, two other sequences of *B. natans* clustered together with kinesin-2 from other rhizarians (**Fig. S1**). Surprisingly, KAP3 was not identified in this species (**Fig. 1B, Table S1**). In conclusion, the kinetoplastid gene configuration with IFT-B proteins, KIN2A and KIN2B, but not KAP, was conserved in all euglenozoans analysed. Since no other kinesin-2 members were identified, we decided to investigate more closely how KIN2A and KIN2B function both *in vitro* and *in vivo*.

**Figure 1:**
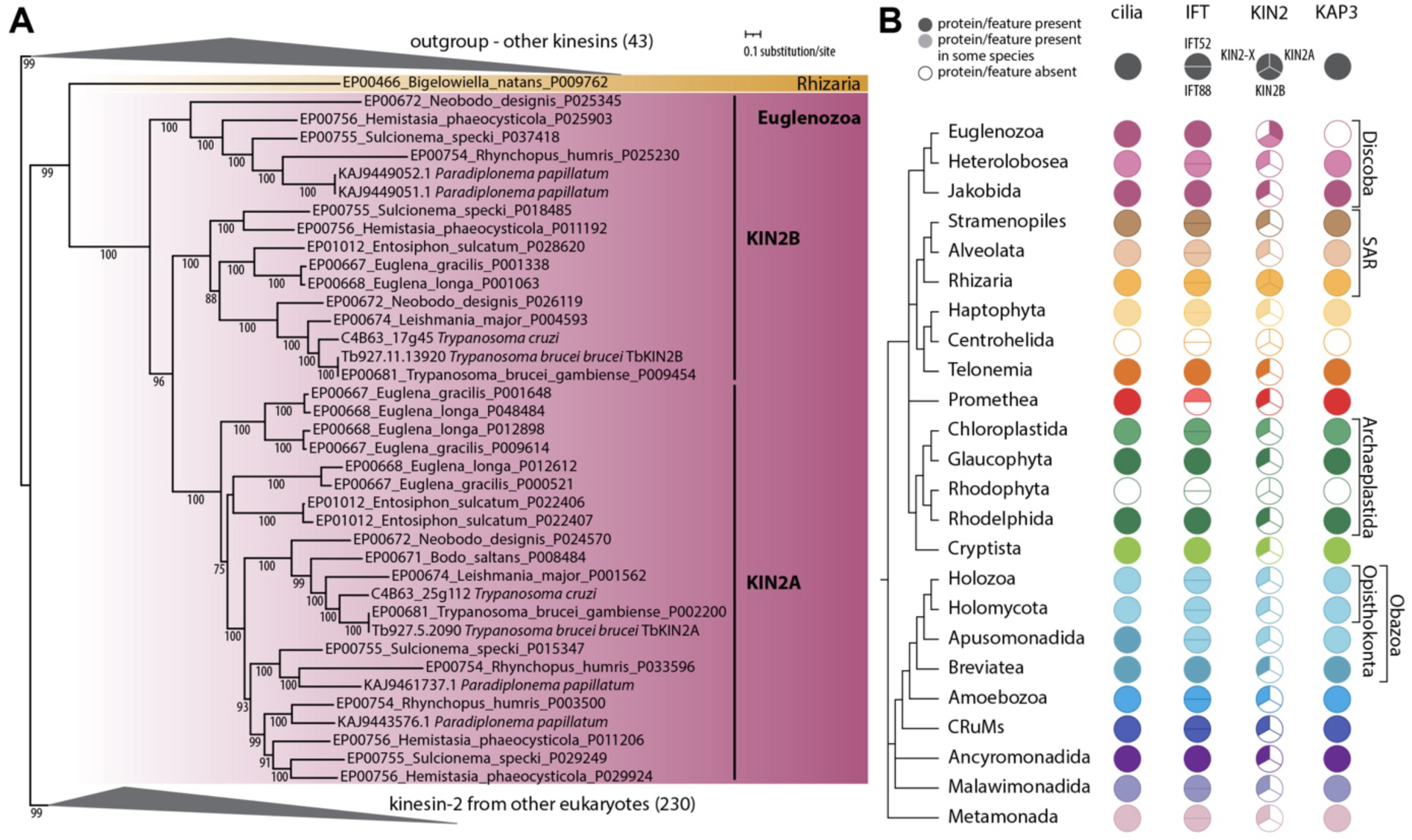
Kinesin-2 and IFT proteins in eukaryotes. **(A)** Phylogenetic analysis of kinesin-2. The maximum-likelihood phylogenetic tree was performed in IQ-TREE using PMSF analysis with 1,000 replicates for ultrafast bootstraps (UFB). Only UFB supports ≥75 are shown. All clades, except for KIN2A and KIN2B, were collapsed for visualisation purposes (for the full tree, see **Fig. S1**). Number in parentheses indicates number of species in the collapsed clade. **(B)** Presence of cilia, IFT proteins, kinesin-2, and KAP in eukaryotes (for details about species, see **Table S1**). Relationships among eukaryotic groups, which are colour-coded, are shown on the left by a schematic tree based on the highly supported relations in the newest studies [86, 112].

### KIN2A and KIN2B can only form homodimers *in vitro*

To investigate the function of KIN2A and KIN2B, we used *T. brucei* as a model system, where IFT has been exhaustively characterised [38, 39]. *T. brucei* KIN2A and KIN2B were C-terminally tagged with HALO and FLAG or SNAP and 6xHis tags, respectively, and expressed *in vitro* in insect cells. FLAG and 6xHis tags were used for protein purification as described previously, while HALO- and SNAP-tags enabled fluorescent labelling for live imaging [40]. KIN2A displayed processive movement on surface-attached mammalian brain microtubules, used here in a standard *in vitro* assay, displaying a slow (0.8 µm/s; **Video 1**, yellow arrow) and a fast (2.4 µm/s; **Video 1**, white arrows) velocity distribution (**Fig. 2A**). The first peak suggests a possible regulation mechanism by the C-terminal tail domain in the absence of cargo in the *in vitro* reconstituted transport assays, as observed previously with the heterodimeric kinesin-2 motor from *C. reinhardtii* [41]. To test this, the C-terminal tail was removed, and this truncated motor faithfully reproduced the fast velocity distribution of the full-length KIN2A motor (2.6 µm/s; **Fig. 2B** and **Video 1**, right panel). To finally demonstrate that the fast velocity distribution represents the fully active state of the KIN2A motor, we completely removed the predicted coiled-coil and random coil domains (i.e. any potential inhibitory domain) and dimerized the catalytical head domains with the coiled-coil domains of the transcription factor GCN4 [42]. This artificially dimerized KIN2A::GCN4 reproduced the fast velocity distribution of the full-length and the C-terminally truncated motors, respectively (2.6 µm/s; **Figs. 2A-C** and **Video 1**, lower panel). Just as KIN2A, the KIN2B motor also moved processively on surface-attached microtubules (**Video 2**) but displayed a single velocity distribution that was comparatively slower than that of the KIN2A motor (**Fig. 2D**). To test if the velocity of the full-length KIN2B represents the fully active state of the motor, we again removed all potentially inhibitory coiled-coil and random coil domains and artificially dimerized catalytical head domains using GCN4. Removal of the entire C-terminus increased the velocity of the motor to 1.9 µm/s (b, **Video 2**, **Fig. 2E**), suggesting that the speed of the full-length KIN2B motor is also partially inhibited by its C-terminus. These results demonstrate for the first time that KIN2A and KIN2B are *bona fide* processive molecular motors that can function by themselves, suggesting that they act as homodimers and that their motility can be at least partially regulated by their C-terminal ends.

**Figure 2:**
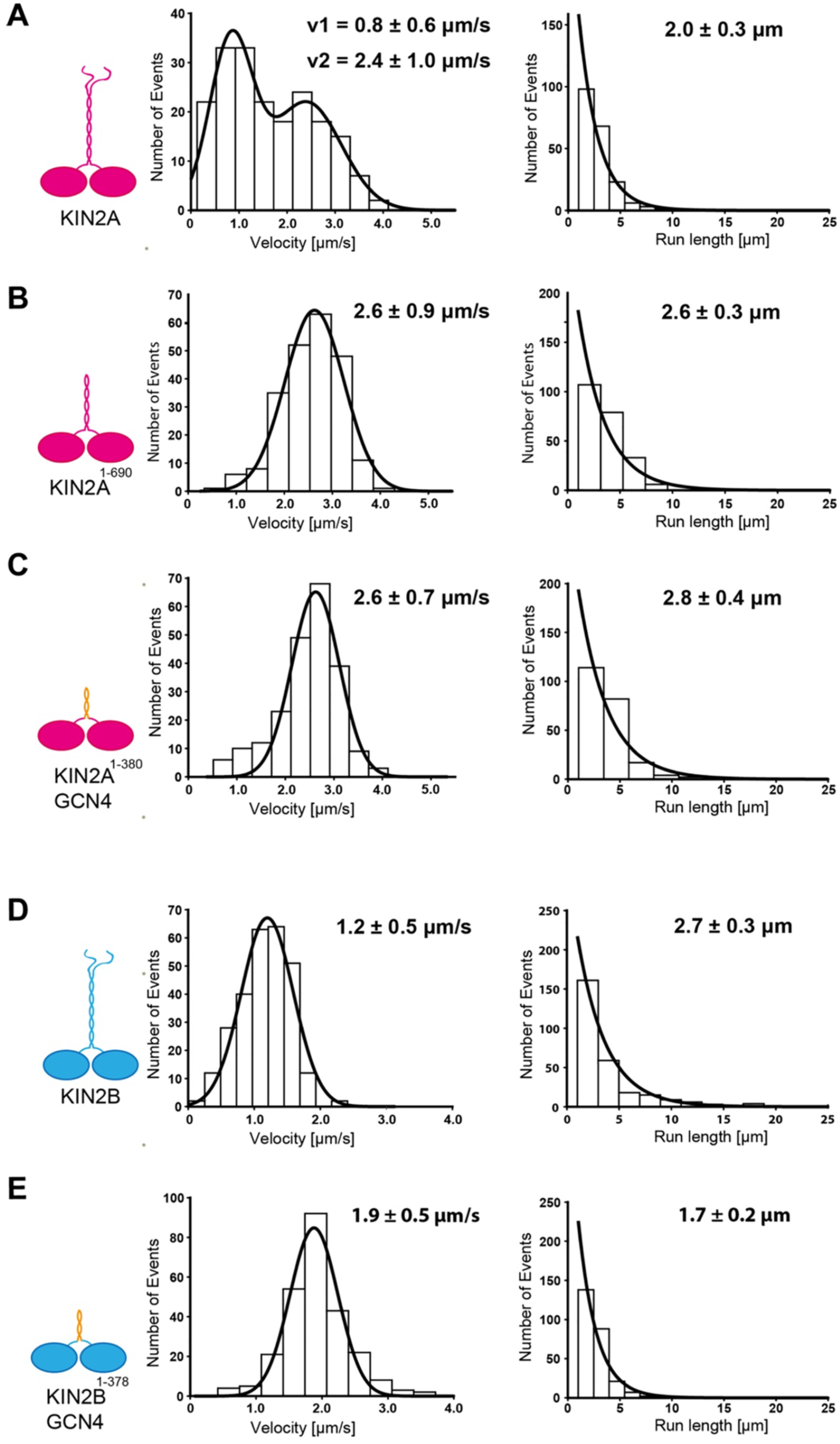
KIN2A and KIN2B from *T. brucei* are processive motors. **(A)** The full length KIN2A motor moved processively on surface-attached microtubules and displayed two distinct velocity distributions, v1 and v2 (n = 194). The slow velocity (v1) indicates that a subpopulation of the wild-type motor might be regulated *via* its C-terminus as observed previously with other kinesin motors. **(B)** Removal of the C-terminus still allowed dimerization (see Fig. 3C) but eliminated the slow velocity (v1) and reproduced the fast velocity distribution (v2) of the full-length KIN2A (n = 225). **(C)** To ensure that the fast velocity distribution (v2) observed with the full-length **(A)** and C-terminally truncated motor **(B)** represents the uninhibited velocity, we artificially dimerized the motor domains (KIN2A^1-380^) with the GCN4 molecular zipper, thus excluding potentially inhibitory domains altogether. This artificially dimerized motor construct also reproduced the fast velocity distribution (v2) of the full-length motor **(A)** as did the C-terminally truncated motor construct **(B)** (n = 219). Contrary to the velocity (A-C, left panels), the run length of the KIN2A motors was not substantially altered by the C-terminal truncations (A-C, right panels). **(D)** The full-length KIN2B moved processively on surface-attached microtubules (n = 246), and **(E)** the velocity increased significantly when the motor domains (KIN2B^1-378^) were dimerized artificially using GCN4 (n = 246). While removal of the C-terminus increased the motor’s velocity, the run length of KIN2B decreased markedly (**E** vs **D**, right panels). The velocity (A-E, left panels) and the run length (**A-E**, right panels) data were obtained from Gaussian (± width of distribution) distribution and single exponential (± confidence interval) fits, respectively. Data were obtained from three independent protein preparations each.

To further corroborate the dimerization states of KIN2A and KIN2B motors, size-exclusion chromatography with multi-angle light scattering (SEC-MALS) was performed. It revealed molecular masses of 252 kDa and 280 kDa for recombinant full-length KIN2A and KIN2B (both FLAG-tagged), respectively, very close to the expected values for homodimers of 248 and 252 kDa (**Fig. 3A-B**). Truncation of the C-terminal tail of KIN2A did not alter its ability to form a dimer (**Fig. 3C**). These data demonstrate that KIN2A and KIN2B form homodimers, but they do not rule out the possibility of a heterodimer formation between KIN2A and KIN2B. To evaluate this possibility, KIN2A fused to a FLAG tag and KIN2B fused to the SNAP-6XHis tag were co-expressed in Sf9 insect cells. A pull-down experiment was performed using an anti-FLAG antibody but only KIN2A::FLAG was precipitated (**Fig. 3D**, lane a). Analysis of the lysate confirmed that sufficient KIN2B::SNAP-6xHis was present (**Fig. 3D**, lane b), suggesting that KIN2A and KIN2B do not form heterodimers at least *in vitro*. By spreading co-expressed fluorescently labelled purified proteins on glass, we next asked how often KIN2A and KIN2B both FLAG-tagged and fluorescently labelled via SNAP and HALO- tags co-localised. As a positive control, we co-expressed the motor subunits KLP11 and KLP20 of the *C. elegans* heterodimeric kinesin-2 [43]. These display exhaustive co-localisation, as expected (**Fig. 3E, top panels**). By contrast, the trypanosome KIN2A and KIN2B did not show association and were stochastically and independently distributed on the sample (**Fig. 3E, bottom panels**). We conclude that the *T. brucei* KIN2A and KIN2B function as homodimers and are not able to constitute a heterodimer, at least *in vitro* and in the absence of any unknown additional subunits.

**Figure 3:**
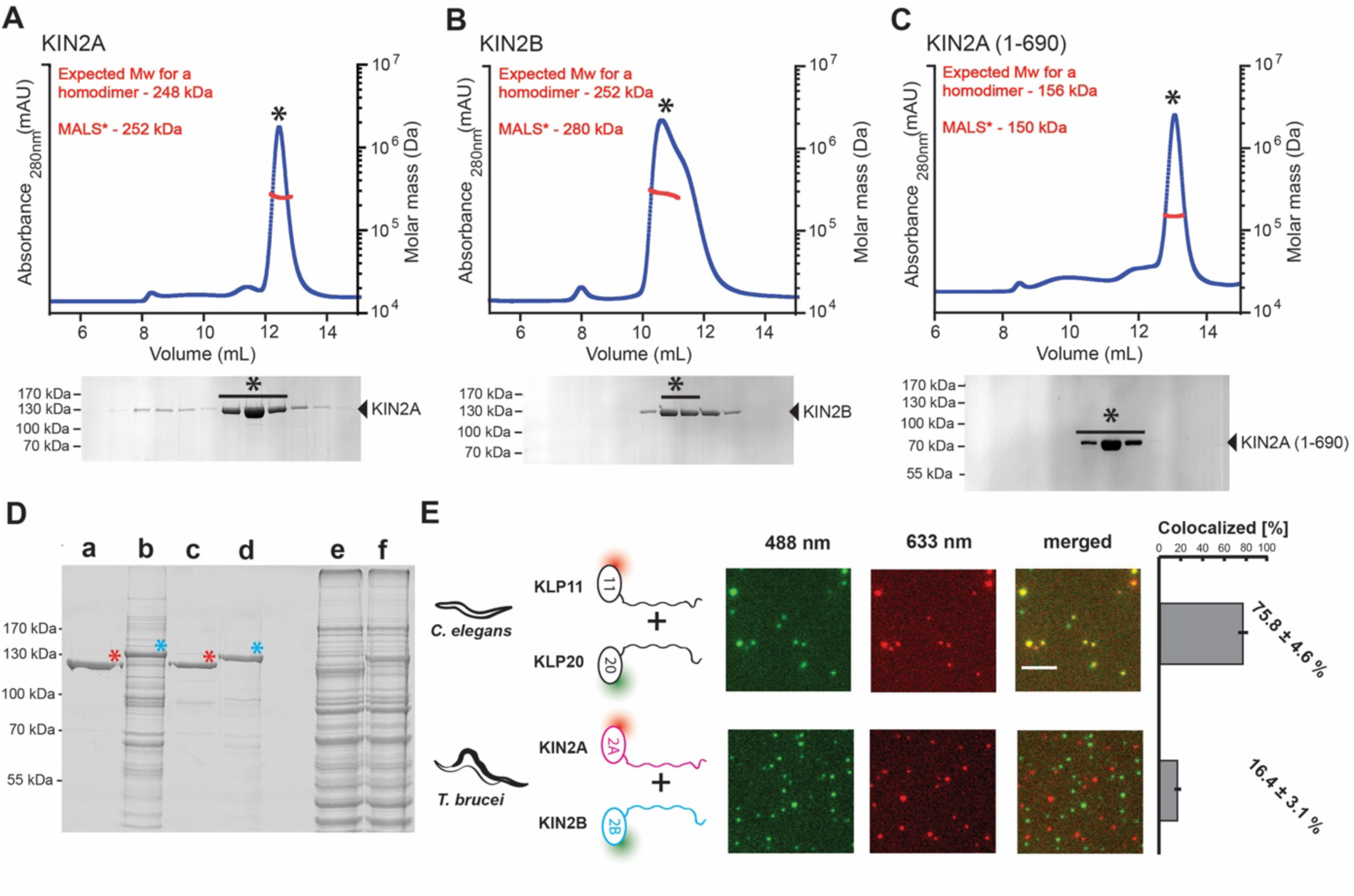
KIN2A and KIN2B from *T. brucei* are homodimeric kinesin-2 motors. (A, B) Size Exclusion Chromatography coupled to Multiple Angle Light Scattering (SEC-MALS) analyses of the recombinantly expressed full-length KIN2A **(A)** and KIN2B **(B)** motors. The respective molecular weights obtained from the MALS fits (red) to the elution profiles (blue traces) are consistent with homodimeric motor proteins (N=3). The bottom panels show the SDS-PAGE analysis of the protein found in the elution peak (asterisk indicates the fractions from which the MALS values were obtained). **(C)** Removal of the C-terminus did not affect the homodimerization of the KIN2A kinesin-2 motor. The truncated KIN2A (1-690) displayed molecular weights that are consistent with homodimeric motor proteins as judged from the SEC-MALS analysis (N=2). The bottom panels show the SDS-PAGE analysis of the protein found in the elution peak (asterisk indicates the fractions from which the MALS values were obtained). **(D)** Flag-tagged KIN2A fails to co-immunoprecipitate KIN2B in co-expression assay and vice versa. Differentially tagged KIN2A^Flag^ and KIN2B^Snap,6xHis^ proteins were co-expressed in Sf9 cells to assess their ability to interact with each other. The Flag-tagged KIN2A was not able to pull down the 6xHis-tagged KIN2B as judged from the SDS-PAGE analysis (lane a, red asterisk). The lysate after the Flag affinity purification was subjected to a His-tag affinity purification to ensure that KIN2B was expressed in sufficient amounts (lane b, blue asterisk). Similarly, Flag-tagged KIN2B did not pull down 6xHis-tagged snap KIN2A as seen in lane c (red asterisk). In addition, the lysate following Flag affinity purification was subjected to a His-tag affinity purification to check whether KIN2A^Snap,6xHis^ was expressed in appropriate amounts (lane d, blue asterisk). Total lysates for each experiment are shown at lanes e and f. **(E)** Co-localization assays with the heterodimeric KLP11/20 from *C. elegans* or the KIN2A and KIN2B motors from *T. brucei* (cartoons are not to scale). The top panel shows the co-localization behaviour of the KLP11/KLP20 motor. As expected from a robust heterodimeric kinesin-2 motor, the SNAP647-labeled KLP11 (633 nm channel) co-localized efficiently with Halo488-labeled KLP20 (488 nm channel) (75.8 ± 4.6%). In contrast, the KIN2A^HALO660^ (633 nm channel) failed to co-localize significantly with KIN2B^SNAP488^ (488 nm channel) (bottom panel). Error bars indicate the standard deviation from three independent protein preparations with ten images each. Scale bar: 5 µm.

### KIN2A and KIN2B display differential localisation along the flagellum

To determine the behaviour of KIN2A and KIN2B *in vivo*, they were tagged endogenously [44] and expressed as fusion proteins with mNeonGreen (mNG) and 6 copies of the Ty-1 tag [45] in *T. brucei* procyclic form cells. The well-characterised IFT-B protein IFT81 [38] was used as a control. Previous proteomic studies showed that KIN2A and KIN2B have low abundance in purified flagella [46], so both alleles (*T. brucei* is diploid) were tagged to guarantee all the protein products were visualised, while only one allele of IFT81 was tagged. To confirm the expression of the fused proteins, we performed a western blot using BB2, an antibody against the Ty-1 epitope (**Fig. 4A**). As expected, the apparent molecular mass of mNG::KIN2A and mNG::KIN2B was at approximately 150 kDa, while IFT81::mNG ran at a slightly lower mass (around 140 kDa), confirming correct integration of the constructs and expression of the fusion proteins. Using an anti-ALBA antibody [47] as a loading control, the relative expression levels were quantified, showing that the KIN2A signal was 39% of the Alba signal and KIN2B 30%, while IFT81 exhibited a similar expression level as Alba (97 and 100%, for the two clones analysed, respectively). Considering only one allele of IFT81 was tagged and assuming that the untagged allele is expressed at the same level as the tagged one, KIN2A and KIN2B have an expression level of 19.5% and 15% of IFT81, respectively, meaning the ratio of IFT81 per KIN2A or KIN2B is approximately between 5:1 to 7:1. We conclude that kinesins are sub-stoichiometric to IFT-B, in line with proteomic analysis of purified flagella from *Chlamydomonas* that indicates a ratio of 8:1 [48].

**Figure 4:**
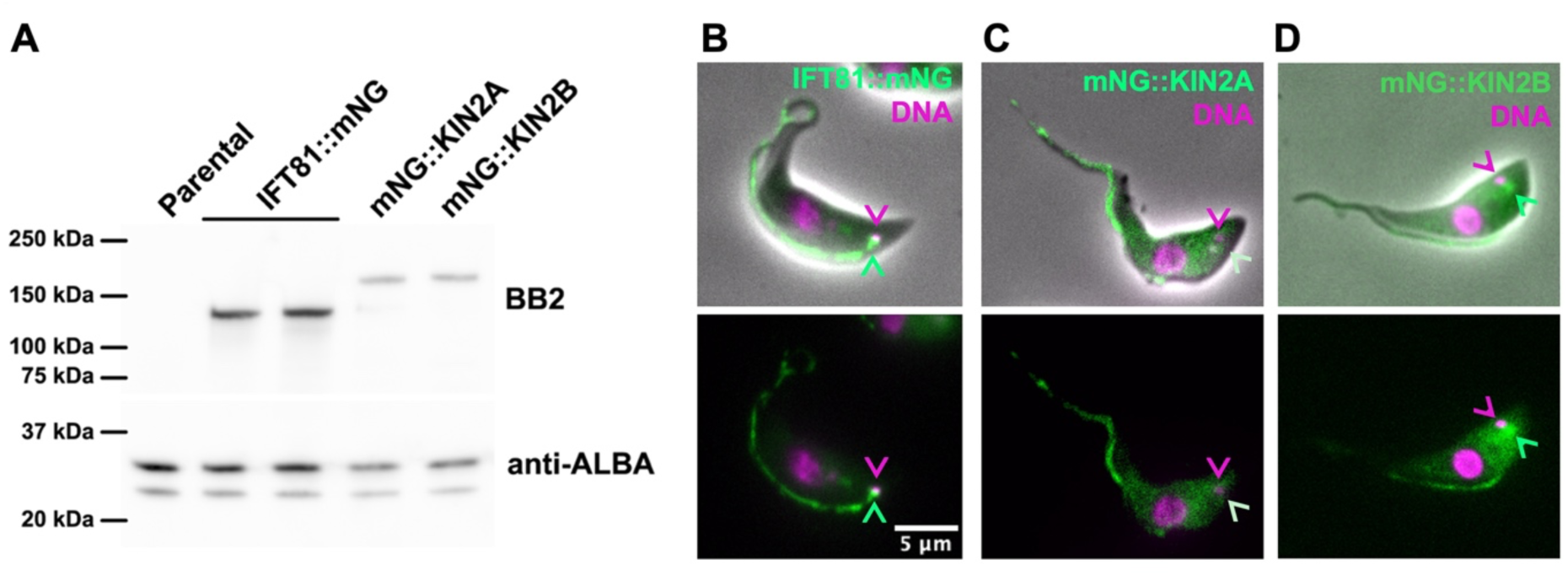
KIN2A and KIN2B localise to the flagellum. The amounts of IFT kinesins were assessed in trypanosomes expressing IFT81::mNG (single allele tagging), mNG::KIN2A, and mNG::KIN2B (both double allele-tagging). **(A)** Western blot analysis shows the protein levels of KIN2A and KIN2B relative to IFT81. The BB2 antibody was used to detect the 6 Ty-1 epitopes in the fusion proteins, with ALBA as the loading control. **(B-D)** Live cells expressing the indicated fusion proteins (green): either IFT81::mNG **(B)**, mNG::KIN2A **(C)** or mNG::KIN2B **(D)** were stained with DAPI (magenta), and images were acquired immediately afterwards. The magenta arrowheads indicate the position of the mitochondrial DNA, which is adjacent to the base of the flagellum [113]. Light green arrowheads point at abundant IFT81::mNG or mNG::KIN2B signal at the base of the flagellum, while the absence of mNG::KIN2A concentration is highlighted by the white arrowhead. Scale bar: 5 µm.

The localisation of the two kinesins was first evaluated by live imaging (**Fig. 4B-D and Fig. 5A,C,E)**. As expected, the control IFT81::mNG produces a bright signal at the flagellar base and is found as numerous particles along the flagellum (**Fig. 4B and Fig. 5A**). Both KIN2A and KIN2B localise to the flagellum but show different distribution patterns. KIN2A signal is evenly distributed along the flagellum without much concentration at the base (**Fig. 4C and Fig. 5C**), while the KIN2B signal appears more intense in the proximal portion of the flagellum and shows a clear basal pool (**Fig. 4D and Fig. 5E**). For both kinesins, some cytoplasmic signal was also detected. These localisations are in line with what was observed by the TrypTag genome-wide subcellular protein localisation project [49]. To make sure that expression of the tagged proteins does not have a negative impact on other IFT proteins or other cellular aspects, we monitored the distribution of the native IFT172 (**Fig. S2A**), cell growth (**Fig. S2B**) and flagellum length (**Fig. S2C**), which were comparable to wild-type cells.

**Figure 5:**
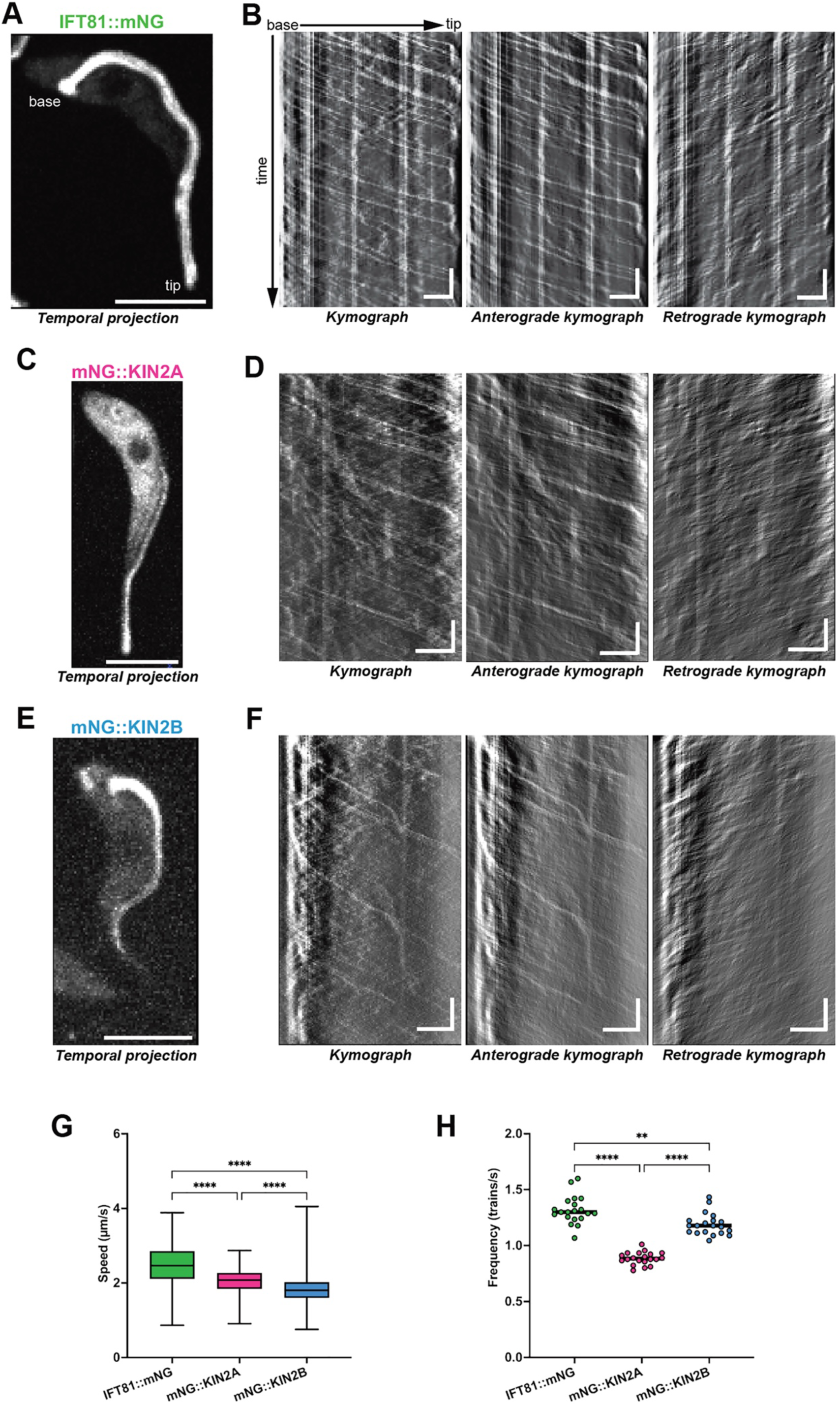
KIN2A and KIN2B cannot independently explain IFT81 anterograde transport. Live-cell imaging was performed on cell lines expressing IFT81::mNG, mNG::KIN2A and mNG::KIN2B. **(A)** Temporal projection of a 30-second acquisition of a single cell expressing IFT81::mNG, showing its concentration at the base of the flagellum and even distribution along the flagellar length. Scale bar: 5 µm. **(B)** Kymographs extracted from the cell shown in A displaying both anterograde and retrograde movement, or with anterograde and retrograde traces extracted separately. Horizontal scale bar: 3 µm; vertical scale bar: 3 s. **(C)** Temporal projection of a cell expressing mNG::KIN2A. Scale bar: 5 µm. **(D)** Kymographs extracted from the cell shown in C displaying both anterograde and retrograde transport. Horizontal scale bar: 3 µm; vertical scale bar: 3 s. **(E)** Temporal projection of a cell expressing mNG::KIN2B, showing a stronger concentration at the proximal portion of the flagellum. Scale bar: 5 µm. **(F)** Kymographs obtained from the cell shown in E. Horizontal scale bar: 3 µm; vertical scale bar: 3 s. **(G)** Speed of the anterograde trains for IFT81::mNG (green), mNG::KIN2A (magenta) and mNG::KIN2B (blue). **(H)** Frequency of the same anterograde trains showing that KIN2A and KIN2B individually do not reach the frequency of IFT81. Data were obtained from three independent experiments, with 20 cells per condition. ****p<0.001, *p=0.0016.

The absence of a KIN2A basal pool was confirmed by immunofluorescence using BB2 to localise KIN2A relative to a transition zone protein, stained with an antibody against FTZC [50], and the axonemal protein SAXO, stained with the mAb25 antibody [51] (**Fig. S2D**). The higher abundance of mNG::KIN2B along the proximal portion of the flagellum is rather unexpected, as none of the 22 IFT proteins investigated to date exhibit such a profile [49, 52–57]. To evaluate if this distribution is correct or a mislocalisation resulting from the fusion to mNG, we expressed a fragment of the *KIN2B* gene as a GST fusion protein in *E. coli*, purified it and injected mice for antibody production. These antibodies were used for indirect immunofluorescence assay (IFA), which revealed that the native protein exhibits the same distribution profile with a strong signal at the base of the flagellum and a higher concentration along the proximal portion of the flagellum in both uniflagellated (**Fig. S3B**) and biflagellated cells (**Fig. S3C**). This signal was similar for all antisera from five immunised mice. To ascertain signal specificity, the IFA experiment was repeated in the *KIN2A2B^RNAi^* cell line [33]. The knockdown triggered a pronounced reduction of the signal in induced cells possessing a short flagellum, demonstrating signal specificity (Fig. **S3D-E**) and arguing against a mis-localisation due to tagging.

As KIN2A and KIN2B are conserved in the Euglenozoa, we investigated their localisation in *Leishmania mexicana* as another trypanosomatid species that can be manipulated in the laboratory. The *L. mexicana* KIN2A and KIN2B were endogenously fused to mNG by CRISPR-Cas9 technology [58]. KIN2A was found throughout the flagellum without clear enrichment at the base (**Fig. S4A,C-E**, top panels) while KIN2B is highly concentrated at the flagellum base and restricted mostly to the proximal portion of the flagellum (**Fig S4B,F-H**, bottom panels). Therefore, the differential distribution of KIN2A and KIN2B is conserved between *Trypanosoma* and *Leishmania*, which are considered at opposite sides of the phylogenetic tree of trypanosomatids [28], suggesting it could be conserved in all members. It should be noted that the KIN2B proximal staining appears proportionally shorter in *L. mexicana* compared to what is found in *T. brucei*, perhaps reflecting different heterogeneities along flagellum length [59, 60]. The flagellar localisation patterns of KIN2A and KIN2B in two different organisms suggest that both kinesins are involved in IFT, aligning with previous data showing that their combined knockdown impacts IFT trains’ speed and frequency as well as flagellum construction [33].

### KIN2A and KIN2B display intraflagellar movement

To investigate the KIN2A and KIN2B relationship with IFT, we performed live-cell imaging on *T. brucei* (**Fig. 5, Fig. S2E-F, Videos 3-5**) and *L. mexicana* **(Fig. S4C-H, Video 6)** tagged cell lines. First, live trypanosomes were observed with confocal spinning-disk microscopy as reported previously [39], and 30-second acquisitions were taken to determine fluorescent particle movement along the flagellum. Using IFT81::mNG as a reference, the temporal projection of one of the videos shows the basal pool and an even distribution of the particles along the flagellum length (**Fig. 5A, Video 3)**. Kymographs were extracted from this acquisition, and both anterograde and retrograde particles were detected (**Fig. 5B**). Since only one allele of IFT81 was labelled, the fusion protein is competing with products of the untagged allele. However, IFT trains are composed of multiple copies of IFT-B complexes with an 8 nm periodicity [61] and their average length of ∼200 nm [38] means that each train should contain at least 25 copies, making it highly likely that the vast majority of trains is labelled. With the same acquisition parameters, mNG::KIN2A particles were also distributed evenly along the flagellar length, but no basal pool was detected (**Fig. 5C, Fig. S2E, Video 4**) as observed previously (**Fig. 4 and Fig. S2**). In agreement with the relatively lower total amount of KIN2A, its flagellar signal looked less intense compared to IFT81::mNG and a notable signal was present in the cell body, except for the nucleus (**Fig. 5C, Fig. S2E**). When the kymographs were extracted, KIN2A particles also displayed anterograde and retrograde movements, indicating their involvement in IFT (**Fig. 5D, Fig. S2F, Video 4**). The mNG::KIN2B temporal projection showed the same pattern as observed before by IFA, with a strong signal at the basal pool and the flagellar proximal portion, but with a weaker signal towards the distal part (**Fig. 5E, Video 5**). The kymographs extracted from this cell confirmed the IFT-like displacement of KIN2B particles, but mostly in the proximal portion of the flagellum, with only a few particles reaching the distal part of the flagellum (**Fig. 5F** arrowheads). The retrograde transport exhibited by both kinesins suggests they are recycled back to the base as cargoes of retrograde trains, as observed in *C. elegans* [21]. Similar conclusions were reached when observing live *L. mexicana* cells expressing fluorescent mNG::KIN2A and mNG::KIN2B. KIN2A was not detected at the base of the flagellum, but robust anterograde and retrograde trafficking was visible (**Video 6,** left panels), as revealed by kymograph extraction (**Fig. S4C-E**). For KIN2B, a strong signal was present at the flagellum base, and as observed previously, the flagellum signal was limited to a shorter portion compared to *T. brucei*. Anterograde and retrograde movement is mostly detected in the very proximal part of the flagellum, with a few occasional particles trafficking in the distal portion (**Video 6,** right panels and **Fig. S4F-H**, bottom panels).

To get a more quantitative view of the contribution of each kinesin to IFT in *T. brucei*, the anterograde particles observed in the three cell lines upon kymograph extraction were analysed, and their speed was measured. This showed that KIN2A and KIN2B particles have a speed compatible with IFT, although they are both a bit slower than IFT81, a difference that is statistically significant (**Fig. 5G**). KIN2B shows a broader speed range and is slower than KIN2A, in agreement with the *in vitro* data (**Fig. 2**). The slightly reduced speed observed for mNG::KIN2A and mNG::KIN2B could be explained by the impact of a fusion reporter protein, although this impacted neither IFT172 distribution, cell growth nor flagellar length (**Fig. S2B-D**). The frequency of anterograde particles was quantified (**Fig. 5H**). Both KIN2A (0.88 ± 0.19 trains/s) and KIN2B (1.18 ± 0.18 trains/s) particles show lower frequency than IFT81 (1.31 ± 0.21 trains/s), indicating that they are unlikely to be present in every anterograde IFT particle. This suggests that neither KIN2A nor KIN2B alone could perform the whole anterograde transport of IFT complexes. If KIN2A and KIN2B are binding independently to IFT trains, the sum would be too high to explain the frequency of IFT81 trafficking. However, two other options could be considered: either some kinesins do not associate to IFTs or some KIN2A and KIN2B associate together to the same IFT train.

### KIN2B is not often associated with IFT trains

These results, together with the fact that most KIN2B particles do not travel to the distal tip of the flagellum, a feature not observed for all the IFT proteins characterised so far, prompted us to examine closely KIN2B behaviour and its potential association with IFT proteins. To approach this, we attempted to produce cell lines in *T. brucei* and in *L. mexicana* that carry both KIN2A or KIN2B tagged with mNG and an IFT protein tagged with a red fluorescent protein in a way that these two proteins could be followed at the same time by two-colour live-cell imaging [38]. However, producing these cell lines was very challenging, and most attempts failed. The presence of a reporter protein on the kinesin motor and another one on an IFT component of the trains may be incompatible with the assembly of the IFT particles or motor association, making live imaging impossible. This has also been reported while attempting to tag two transition fibre proteins in *T. brucei* [62], suggesting that steric hindrance in a densely packed environment might be an issue. Moreover, the only fluorescent reporter tolerated for KIN2A and KIN2B in single tagging was mNG, forbidding the monitoring of these two motors within the same cell. However, the combination of mNG::KIN2B with endogenously tagged tdT::IFT140 was the only one to result in a viable cell line where both signals were positive and exhibited the same profile as in single tagging. Live-cell imaging was carried out in this cell line using a confocal spinning disk equipped with a beam splitter and two cameras, which allows following these two signals at the same time (**Fig. 6, Video 7**). For the mNG::KIN2B signal, multiple anterograde particles restricted to the proximal portion of the flagellum are identified, but they do not exhibit the tdT::IFT140 signal (**Fig. 6A**). By contrast, only a few KIN2B particles reach the flagellum distal portion, named full-length particles, and these show colocalization with the IFT140 signal (**Fig. 6B, H**). Merging the KIN2B and IFT140 kymographs confirmed that most proximal KIN2B particles did not overlay with IFT140 (**Fig. 6C, F**). The proximal and full-length KIN2B anterograde particles were analysed separately, confirming the visual impression that the full-length particles exhibit a frequency ∼4- fold lower than the proximal ones (**Fig. 6D**) but display faster motility (**Fig. 6E**). The slower speed exhibited by the proximal KIN2B particles with the limited overlay with IFT140 in the anterograde kymographs suggested that they are rarely associated with IFT trains. By contrast, a faster speed of KIN2B particles associated with IFT140 is compatible with *in vitro* data that indicate partial regulation of KIN2B in the absence of the tail (**Fig. 2E**), which usually interacts with IFT proteins. To quantify this, we analysed the mNG::KIN2B and tdT::IFT140 overlay using the Manders overlap coefficient [63, 64]. The mNG::KIN2B tracks used to quantify the particles’ frequency and speed were filtered into proximal (**Fig. 6F**) and full-length (**Fig. 6H**) according to a delimitation of the proximal two-thirds of the flagellar length. These tracks were used as regions of interest, and their signal intensity was compared to the tdT::IFT140 tracks (**Fig. 6, G and I**). The M1 index indicates the percentage of the KIN2B signal that overlaps with the IFT140 signal, while the M2 index exhibits the percentage of the IFT140 total signal that overlays with the KIN2B signal. 36% of the KIN2B proximal particles overlay with IFT140, and only 12% of the IFT140 particles overlap with KIN2B proximal particles (**Fig. 6G**), indicating the KIN2B proximal particles exhibit little or no association with IFT trains. The opposite was observed for the full-length KIN2B particles, which show 83% overlap with the IFT140 signal, with some cells exhibiting complete signal overlap (**Fig. 6I**). 32% of the IFT140 particles overlap with KIN2B full-length particles, matching the average frequency obtained for the full-length particles. This suggests that the KIN2B full-length particles are the ones associated with IFT trains and so are involved in the anterograde transport. The observation of KIN2B proximal particles not being associated with IFT trains discarded the hypothesis of KIN2A and KIN2B cooperating on a simple handover mechanism at the end of the proximal portion of the flagellum. However, the low frequency of these particles means that about two-thirds of the IFT proteins must be transported by a different motor, presumably KIN2A. Indeed, the combination of KIN2A (0.88 ± 0.19 trains/s) and KIN2B full-length (0.43 ± 0.08 trains/s) particles’ frequencies results in the exact frequency observed for IFT81 (1.31 ± 0.21 trains/s) or IFT140 (1.34 ± 0.13 trains/s), suggesting these motors carry different IFT anterograde trains and function as independent homodimers *in vivo*, as suggested from *in vitro* data.

**Figure 6:**
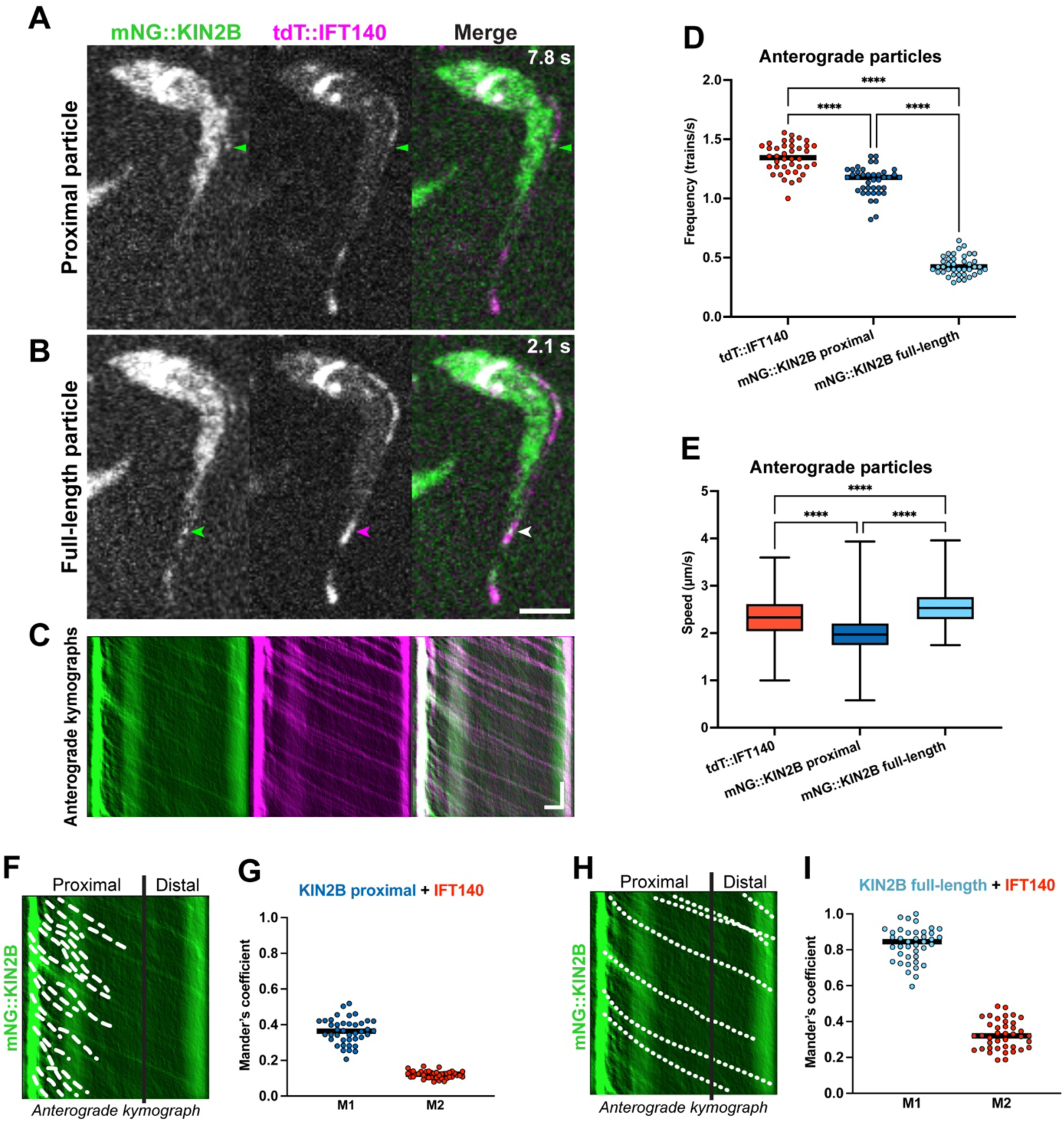
KIN2B is not always associated with IFT trains. IFT was analysed in a cell line expressing mNG::KIN2B and tdT::IFT140, with the displacement of these two proteins followed simultaneously in live cells. **(A)** A single frame from a 15-second acquisition shows a proximal mNG::KIN2B particle that lacks a tdT::IFT140 signal (green arrowhead). **(B)** Another frame from the same 15-second acquisition as in **A** shows a full-length mNG::KIN2B particle (green arrowhead) with a tdT::IFT140 signal (magenta arrowhead, white on the merged panel). Scale bar: 5 µm. **(C)** Anterograde kymographs extracted from the acquisition shown in **A** and **B**. Horizontal scale bar: 3 µm; vertical scale bar: 3 s. **(D)** Frequency of anterograde trains in the double-tagged cell line, with IFT140 (red) and KIN2B particles categorised as proximal (dark blue) or full-length (light blue). ****p<0.001. **(E)** Speed of the anterograde particles, obtained from 15-second acquisitions of 20 cells in each of three independent experiments. ****p<0.001. **(F)** The mNG::KIN2B anterograde kymograph from **C**, with the proximal portion of the flagellum delimited and tracks annotated as proximal particles. **(G)** Manders’ coefficient of colocalisation between mNG::KIN2B proximal particles and tdT::IFT140 particles. M1 represents the rate of overlap between the mNG::KIN2B proximal signal and the tdT::IFT140 signal, while M2 represents the rate of overlap between the tdT::IFT140 signal and mNG::KIN2B proximal particles. This analysis was conducted on the particles from **D** and **E**. **(H)** The mNG::KIN2B anterograde kymograph from **C,** with tracks annotated as full-length particles. (**I**) Manders’ coefficient of colocalisation between mNG::KIN2B full-length particles and tdT::IFT140 particles. M1 represents the rate of overlap between the mNG::KIN2B full-length signal and the tdT::IFT140 signal, while M2 represents the rate of overlap between the tdT::IFT140 signal and mNG::KIN2B full-length particles. This analysis was conducted on the particles from **D** and **E**.

### KIN2A is not essential for flagellum construction

As KIN2A and KIN2B seem to cooperate for the anterograde transport of IFT trains but in a distinct mechanism from what is known for organisms with two distinct IFT kinesins, we carried out functional analysis by generating mutants for these two motors. Individual RNAi knockdown did not produce a visible flagellar phenotype in procyclic trypanosomes [33], so we used a CRISPR-Cas9 gene deletion approach [58]. We started with KIN2A, which is the motor responsible for ∼70% of the anterograde transport in wild-type cells. The correct gene deletion was confirmed by PCR (**Fig S5A-B**) and whole-genome sequencing (**Fig S5C**). *KIN2A-*ablated cells were viable and exhibited an apparent normal morphology, as shown for example by tubulin and DNA staining (**Fig. 7A**). They grew a bit more slowly than parental cells (**Fig. 7B**) and possessed slightly shorter flagella (17.7 ± 3.2 µm in *KIN2A^-/-^* vs 19.1 ± 3.1 µm on parental cells) (**Fig. 7C**). A possible impact on flagellum structure was searched by transmission electron microscopy, but no visible changes were detected (**Fig. S5D**). In *T. brucei*, IFT trains are restricted to the doublet microtubules 3-4 and 7-8 [38, 65] and this was also the case for the *KIN2A*^-/-^ cell line (**Fig. S5E**).

**Figure 7:**
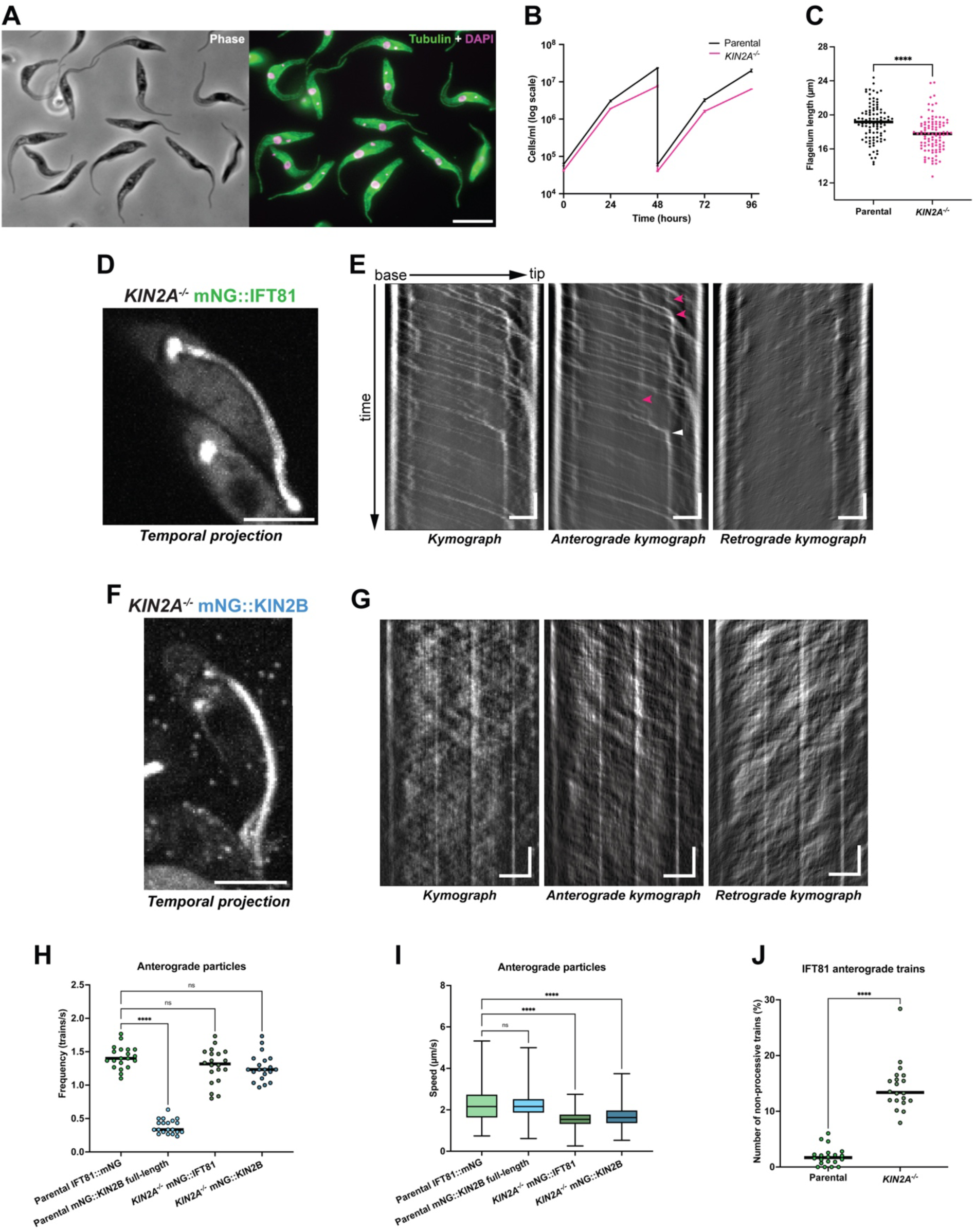
KIN2A deletion has a mild impact on flagellum construction and IFT. **(A)** Representative image of *KIN2A^-/-^* cells stained with an antibody against long tubulin poly-glutamylation chains (polyE, green) and DAPI (magenta). Scale bar: 10 µm. **(B)** Growth curves showing that cells grow slightly slower after *KIN2A* deletion. **(C)** Comparison of flagellar length in parental versus *KIN2A^-/-^* cells (*n* = 100, two independent experiments). ****p<0.001. **(D)** Temporal projection of a 30-second acquisition of a *KIN2A*-deleted cell expressing mNG::IFT81. Scale bar: 5 µm. **(E)** Kymographs extracted from the cell shown in **D**. Some anterograde trains exhibit non-processive behaviour, either changing speed (pink arrowheads) or becoming arrested (white arrowhead) along the flagellum. Horizontal scale bar: 3 µm, vertical scale bar: 3 s. **(F)** Temporal projection of a *KIN2A*-deleted cell expressing mNG::KIN2B, showing its even distribution along the flagellum. Scale bar: 5 µm. **(G)** Kymographs extracted from the cell shown in **F** following background subtraction. Horizontal scale bar: 3 µm, vertical scale bar: 3 s. **(H)** Frequency of anterograde trains in *KIN2A^-/-^* cells expressing mNG::IFT81 (dark green) or mNG::KIN2B (dark blue), compared to parental cells expressing IFT81::mNG (light green) or mNG::KIN2B (light blue, only full-length particles were considered). Data were obtained from 20 cells in each of three independent experiments (except *KIN2A^-/-^* mNG::KIN2B, where only one experiment was conducted). ****p<0.001, ns: not significant. (**I**) Speed of anterograde trains analysed in **H**. ****p<0.001, ns: not significant. **(J)** Comparison of the percentage of non-processive IFT81 trains present in parental versus *KIN2A^-/-^* cells. ****p<0.001.

The impact of *KIN2A* deletion on IFT trafficking was assessed by live-cell imaging following expression of mNG::IFT81 (**Fig. 7D-E, Video 8**). The temporal projection of a 30-second acquisition (**Fig. 7D**) showed that IFT81 distribution along the flagellum is similar to what is observed in wild-type cells (**Fig. 5A**). However, the kymograph extraction indicated that the anterograde particles are less processive upon *KIN2A* deletion (**Fig. 7E**, pink arrowheads), with particles changing speed along the length and sometimes staying arrested before reaching the distal tip (**Fig. 7E**, white arrowhead). Still, the total frequency of these IFT81 particles remained the same as observed before deletion (**Fig. 7H**). However, their average speed was affected, as *KIN2A^-/-^* cells exhibited slower anterograde trafficking (**Fig. 7I**). The percentage of anterograde trains that changed speed or got arrested along the flagellar length was quantified and showed a significant increase rising from 1% in control cells to 14% following *KIN2A* deletion (**Fig. 7J**).

Although KIN2A deletion had an impact on IFT, this relatively mild phenotype was unexpected as KIN2A is responsible for most of the anterograde transport in control cells. This suggests that KIN2B can compensate for the absence of KIN2A. This hypothesis was confirmed by the expression of mNG::KIN2B in *KIN2A^-/-^* cells (**Fig. 7F-G, Video 9**). The temporal projection showed that *KIN2A* deletion leads to KIN2B presence throughout the flagellum (**Fig. 7F, Video 9**), which is in stark contrast with its concentration in the proximal portion of the flagellum observed in control cells (**Fig. 5E**). Moreover, the concentration of mNG::KIN2B at the flagellar base appears less striking. Such a bright signal poses a challenge for IFT analysis, masking most trafficking events (**Video 9**). This problem was partially overcome by performing background subtraction on the kymographs extracted from these acquisitions, which revealed that most KIN2B particles were now trafficking through the whole length of the flagellum (**Fig. 7G**). The frequency analysis revealed that KIN2B full-length particles are much more abundant in *KIN2A^-/-^*cells than those observed in wild-type cells and are now equally abundant as IFT81 (**Fig. 7H**). This suggests that KIN2B is carrying all IFT anterograde trains after KIN2A depletion. KIN2B and IFT81 particles also display slower speeds compared to control cells (**Fig. 7I**), supporting this hypothesis.

Next, the well-established CRISPR-Cas9 strategy [58] was used to individually delete both copies of *L. mexicana KIN2A* genes, as confirmed by PCR analysis (**Fig. S6A**). Morphological analysis revealed that *L. mexicana KIN2A^-/-^* parasites looked essentially normal, except for their flagella which were shorter compared to control cells (12.6 ± 2.6 µm, n=54 instead of 17.7 ± 4.6 µm, n=64). This is similar to what was observed for trypanosomes, except that it is a bit more pronounced, perhaps reflecting greater flexibility in flagellum length in *L. mexicana* [66] compared to *T. brucei* [33]. Examination by transmission electron microscopy showed that the overall cytological architecture looked normal, including the flagellum (**Fig. S6B**). These results therefore phenocopy the deletion of KIN2A in *T. brucei* and confirm that KIN2A is not essential for flagellum construction, despite being the most extensively used motor for anterograde IFT trains in the wild-type situation.

### KIN2B is critical for flagellum construction

KIN2A absence is compensated by KIN2B, explaining its limited impact on flagellum construction and suggesting that these two kinesins are functionally redundant. To challenge this hypothesis, we moved forward with *KIN2B* deletion. Although KIN2B contributes only 30% of the anterograde transport, its deletion turned out to be non-viable in *T. brucei* despite multiple rigorous attempts. This suggests that KIN2A cannot compensate for the loss of KIN2B and argues against redundancy. We therefore performed a double knockout of KIN2B in *L. mexicana* since, unlike *T. brucei*, it can proliferate in culture without its flagellum, albeit more slowly [67–69]. Deletion of *KIN2B* was confirmed by PCR analysis (**Fig. S6C**) and yielded a clear phenotype with cells failing to assemble a normal flagellum. Ultrastructural analysis revealed that only the basal body and a short transition zone were visible, with the axoneme being absent (**Fig. 8A**). In a few cases (8.3% of the cells), a tiny membrane stub was visible, but this does not contain either microtubule doublets or other structural elements (**Fig. 8A**). This phenotype is similar to the one obtained upon deletion of other IFT components [67, 69] and demonstrates that KIN2B is essential for flagellum construction in *L. mexicana*. This explains the failure in obtaining the double knockout in *T. brucei* since the flagellum is essential in this organism, even in culture, due to its contribution to morphogenesis during cell division [52, 56, 70, 71]. To evaluate how KIN2B, which transports only a minority of IFT trains, could be essential for flagellum construction, the distribution of two well-characterised IFT proteins was examined upon *in situ* tagging with mNG::IFT140 as marker of the IFT-A complex (**Fig. 8B**) and IFT172 as representative of the IFT-B complex (**Fig. 8C**). While both proteins were detected at the base and along the flagellum in control cells where they displayed regular IFT, they appeared highly concentrated in the basal body region in the *KIN2B^-/-^* cell line (**Fig. 8B-C**), which can easily be determined thanks to their proximity to the kinetoplast (mitochondrial genome) attached to the basal body. These results demonstrate that KIN2B is essential for flagellum construction but not for targeting of IFT proteins to the basal body area.

**Figure 8:**
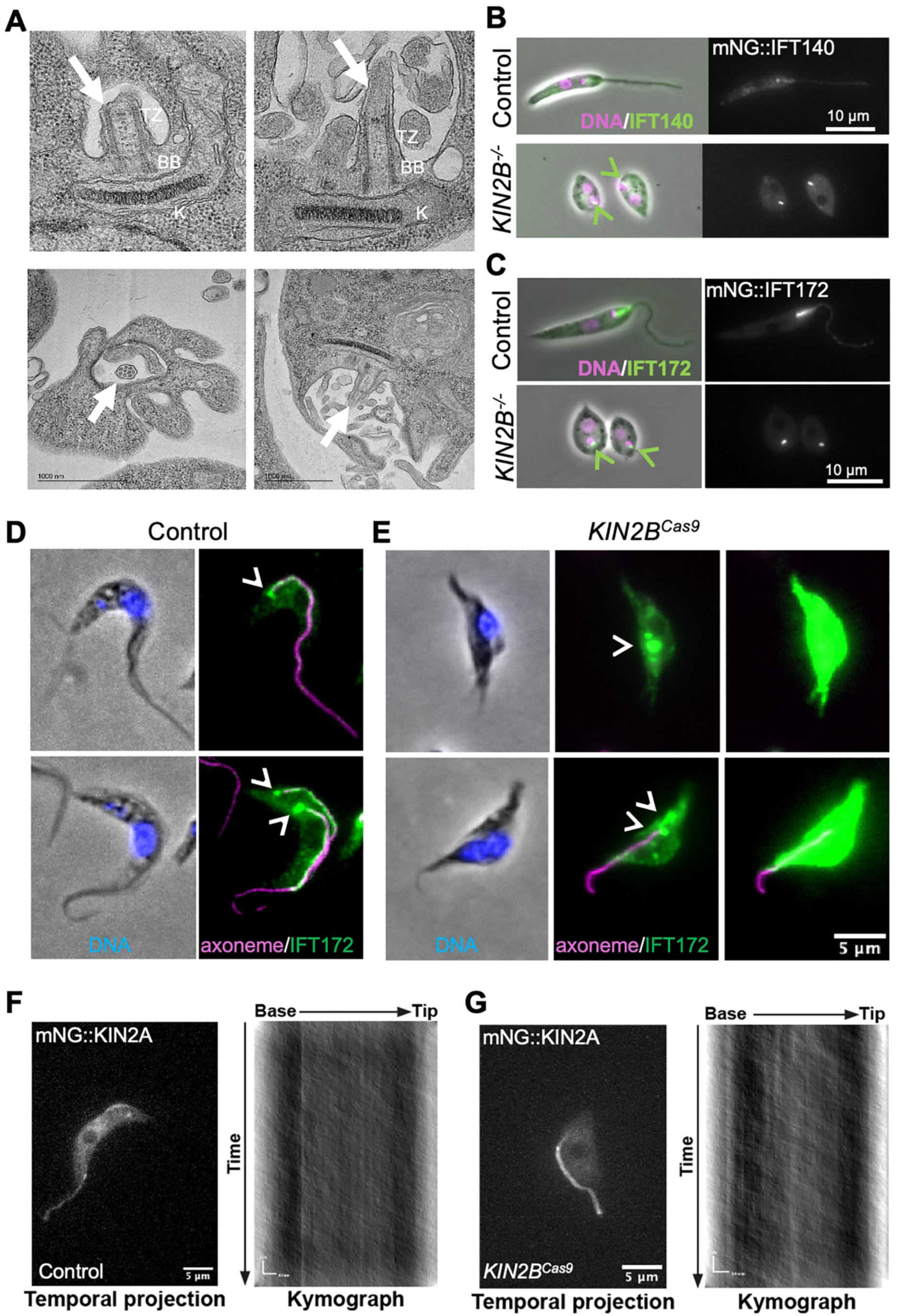
KIN2B is essential for flagellum construction. **(A)** Sections through the flagellar pocket area of *L. mexicana KIN2B^-/-^* cells revealing the presence of a tiny flagellum (arrow) composed of a short transition zone (TZ) and an apparently normal basal body (BB). K, kinetoplast (mitochondrial genome). **(B,C)** The *L. mexicana KIN2B^-/-^* cell line was transformed to express either mNG::IFT140 **(B)** or mNG::IFT172 **(C)** as representative of the IFT-A and the IFT-B complex, respectively. The top panels show an example of a parental cell with its flagellum. The signal for mNG::IFT140 **(B)** and mNG::IFT172 **(C)**(both in green) is pronounced at the base and at the proximal part of the flagellum and more discrete along the length of the axoneme. The two bottom panels show representative examples of *KIN2B^-/-^* cells where the fusion proteins are found in a bright spot close to the kinetoplast (green arrowheads), in agreement with a localisation at the base of the tiny flagellum. Cells were stained with DAPI (magenta) to reveal nuclear and mitochondrial genomes. **(D,E)** Trypanosomes expressing mNG::KIN2A were nucleofected with a mixture containing Cas9, gRNA and repair templates to introduce stop codons in each reading frame of the *KIN2B* gene (*KIN2B^Cas9^*). Cells were grown for 48h, fixed and processed for IFA with mAb25 (marker of the axoneme, magenta), the antibody against IFT172 (green) and DAPI (blue). **(D)** Representative examples of control cells, showing the typical IFT staining at the base (arrowheads) and along each flagellum in a uniflagellated (top) or a biflagellated (bottom) cell. **(E)** Representative examples of *KIN2B* Cas9-inactivated cells, showing a non-flagellated trypanosome with a massive accumulation of IFT172 (arrowheads) in the basal body area (top) and a cell with a short old flagellum that duplicated its basal bodies (arrowheads) but failed to elongate a new flagellum (bottom). The central images were optimised to show the IFT172 profile, while the right image was normalised against the control and is saturated, indicating accumulation of IFT172 throughout the cell body. **(F-G)** KIN2B disruption does not impact mNG::KIN2A trafficking in *T. brucei*. In control cells **(F)**, mNG::KIN2A circulates along the whole length of the flagellum. **(G)** When KIN2B is disrupted (*KIN2B^Cas9^*), trypanosomes that retain a short old flagellum (where the amount of KIN2B had been reduced in the previous generation), mNG::KIN2A still circulates normally. Left panels show the temporal projections and right panels show the corresponding kymographs.

We tried to tag the *Leishmania* KIN2A to monitor its fate in the absence of KIN2B, but despite several attempts, the cell lines either did not grow, or the fluorescent protein expression was limited to a small sub-population of cells. Ideally, an inducible RNAi knockdown strategy would allow monitoring the impact of KIN2B depletion on IFT and KIN2A in *T. brucei* over time, as performed for multiple IFT proteins [52]. Unfortunately, RNAi turned out to be ineffective in silencing KIN2B individually in procyclic cells [33]. We therefore reconsidered how to inactivate KIN2B in *T. brucei* and moved to another strategy recently developed in the field, consisting of transfecting the Cas9 enzyme combined with guide RNAs (gRNAs) and a repair template, which can achieve high efficiency in the absence of drug selection markers (up to 60% of cells with a fluorescent reporter)[72]. To demonstrate that this approach can be used to study IFT-related genes, stop codons were introduced in all three reading frames of the *IFT172* gene and the impact of Cas9 targeting was monitored in trypanosomes (**Fig. S7A-B**). Disrupting *IFT172* led to an arrest in new flagellum formation and emergence of short non-flagellated cells 24 to 48 hours after nucleofection (**Fig. S7B**, white arrows). Cells with shorter flagella are also visible (**Fig. S7B**, yellow arrows), reflecting intermediate stages as a result of protein turnover following gene deletion [33, 71]. This is also exemplified by the presence of trypanosomes that still possessed the old flagellum, duplicated nucleus and kinetoplast but failed to assemble a new flagellum (**Fig. S7B**, orange arrows) as a result of IFT172 gene inactivation. Immunofluorescence with the anti-IFT172 validated a significant drop of IFT172 abundance and absence of signal in all non-flagellated trypanosomes, as well as in many flagellated ones (**Fig. S7**), in agreement with IFT protein turnover in flagella constructed before IFT172 depletion [73]. Despite the presence of normal cells that were presumably not successfully transfected, about half of the population showed reduced or absent IFT172 after 48h, with all such cell profiles phenocopying IFT172 silencing by RNAi knockdown [52].

Having validated the approach, gRNAs and repair templates were designed to introduce stop codons in the three reading frames of the *KIN2B* gene. To evaluate the efficiency of KIN2B depletion, the cell line expressing mNG::KIN2B was used. Western blotting with BB2 showed a pronounced reduction in the total amount of mNG::KIN2B 48h after nucleofection of Cas9 associated with *KIN2B*-gRNA and repair templates (**Fig. S8A**). Co-staining with mAb25 and anti-FTZC as markers of the axoneme and the transition zone, respectively, revealed the emergence of non-flagellated cells as well as others with a clearly shorter flagellum (**Fig. 8D-E, Fig. S8B**). In some non-flagellated cells, the staining for FTZC was reduced or even absent, indicating an impact on formation of the transition zone, something that had been noticed previously in the case of IFT-B knockdowns [74]. These cellular phenotypes are similar to those obtained upon targeting of IFT172 and reveal an essential contribution of KIN2B to flagellum construction, as observed in *L. mexicana*.

To evaluate the impact of KIN2B deletion on IFT protein distribution, these cells were stained by IFA with the anti-IFT172 antibody (**Fig. 8D-E** and **S8B**). In control trypanosomes, IFT172 is found at the base of the flagellum and spreads heterogeneously along its length as expected [52](**Fig. 8D**). By contrast, IFT172 appears highly concentrated at the basal body area in most non-flagellated cells (**Fig. 8E**, top panels and **Fig. S8B**, white arrows). We next examined cells with short flagella since these must have been depleted of KIN2B in the generation before. First, we looked at IFT172 distribution and found out that it is also highly concentrated at the base of the short flagellum and in the cytosol (**Fig. 8E**, bottom panels and **Fig. S8B**, yellow arrows), as observed for non-flagellated trypanosomes. This result suggests that KIN2B is essential for IFT protein import in the flagellum or, possibly, for IFT train formation. Second, we evaluated the impact of KIN2B absence on KIN2A distribution. The Cas9 experiment targeting KIN2B was reproduced in the trypanosome cell line expressing mNG::KIN2A. After 48h, cells with shorter or no flagella emerged, as described above. Strikingly, live imaging showed that mNG::KIN2A traffics normally in short flagella (**Fig. 8F-G**), revealing that KIN2A is still able to access flagella in the absence of KIN2B, in contrast to IFT172 that is limited to the base of the flagellum (**Fig. 8E**). Therefore, KIN2B plays a key role in flagellum construction by ensuring the entry of IFT proteins and/or IFT trains, but not of KIN2A, in the flagellar compartment.

## Discussion

### The presence of two homodimeric kinesins reflects a candidate euglenozoan synapomorphy

The heterotrimeric kinesin-2 is considered the main IFT anterograde motor, being essential for the assembly of cilia and flagella in several organisms, ranging from mammals to unicellular organisms, including the green alga *C. reinhardtii*, where it is actually the only kinesin-2 motor [13, 75]. The absence of KAP and the discovery of two candidate homodimeric kinesins in kinetoplastids [29, 30] therefore raise questions about the origin and the function of these atypical kinesins. The presence of KIN2A and KIN2B and the concomitant absence of KAP in the free-living Euglenida and Diplonemida species rules out a potential specificity to parasitism (for example, in the case of flagellum attachment to host tissues [76]) and favours an alternative explanation. It seems that the KIN2A and KIN2B presence is another euglenozoan synapomorphy, extending the list of common features of these protists (such as polycistronic transcription and spliced leader *trans*-splicing of nuclear transcripts, the presence of sub-pellicular microtubules and a paraflagellar rod, etc.) [27].

Kinetoplastids also single out by having two distinct genes encoding IFT dynein heavy chains [29, 67] and we showed previously that these function as a heterodimer, with each subunit being essential for retrograde transport and for flagellum assembly in *T. brucei* [53]. Intriguingly, structural studies of the *in vitro* assembled human IFT dynein complex revealed that although it is constituted of a homodimer of heavy chains, each of them adopts a distinct configuration to match the repeats of the IFT-B complexes in the IFT train [77]. Therefore, the presence of two different dynein heavy chains might in fact reflect a different strategy to adapt to constraints imposed by the IFT system.

### KIN2A and KIN2B are homodimeric motors with different flagellar localisation and function

The *in vitro* investigations reported here show that KIN2A and KIN2B function as homodimeric motors that are at least partially regulated in the absence of a cargo. This is a well-established feature of motors of the kinesin-2 family, such as the homodimeric OSM-3 from *C. elegans* or KIF17 from mouse [78, 79]. In the case of OSM-3, subunits of the IFT-B complex were identified to relieve the auto-inhibition [80]. In fact, the presence of only two IFT-B subunits is sufficient to reproduce the *in vivo* velocity of the OSM-3 motor in functional *in vitro* reconstitution assays [43]. It is therefore conceivable that the subunits of the IFT-B complex are also involved in the full activation of the homodimeric KIN2A and KIN2B motors in *T. brucei in vivo*.

The *in vivo* situation appears more complex both in *T. brucei* and *L. mexicana*, with KIN2A showing IFT trafficking throughout the length of the flagellum while most of the KIN2B pool is restricted to the base of the flagellum and its proximal portion, with only a limited proportion displaying trafficking from the base to the tip. The localisation of KIN2A had been previously reported in the *T. brucei* bloodstream stage using a rabbit antiserum raised against aa 391-696 of KIN2A, which labeled the flagellum, the basal body area and the cytoplasm [30]. Using the same antiserum provided by Dr. Welch, we confiremd this localisation in procyclic trypanosomes. However, while expression of *KIN2B* double-stranded RNA produced an 8-fold reduction of the signal obtained with this antibody by western blot, it did not impact the IFA signal at the flagellar base (our unpublished data), questioning antibody specificity in this assay. KIN2A turned out to be a faster motor (2.5 µm/s) than KIN2B (1.9 µm/s) during *in vitro* assay, a trend confirmed in live trypanosomes, although with a lower difference, perhaps due to the more complex cellular environment: trafficking on doublets instead of singlets, proximity of the membrane and/or presence of IFT proteins as cargoes. Moreover, kinesin regulators present in the flagellar compartment, such as protein kinases, could alter the behaviour of molecular motors [81]. Live imaging analysis revealed that KIN2A ensures the majority of IFT protein trafficking, hence it was unexpected to discover that KIN2A is dispensable to assemble flagella while KIN2B is essential in both *T. brucei* and *L. mexicana*. This explains why the null mutant could not be obtained in *T. brucei* since its flagellum is required for cell growth in this organism, even in culture [71]. These results differ from the RNAi knockdown results published on the bloodstream stage of the parasite where knockdown of KIN2A, but not KIN2B, turned out to be lethal. This could be explained by the less potent efficiency of RNAi against KIN2B [30]. Since bloodstream cells are more sensitive to flagellar perturbations [34, 35], the reduced flagellar length observed here upon deletion of KIN2A might be sufficient to interfere with cell division.

### KIN2A and KIN2B could function in a division-of-labour model

To integrate all these data, we propose a division-of-labour model (**Fig. 9**, top panel) where KIN2B (blue) is responsible for the delivery of IFT proteins in the flagellar compartment while KIN2A (magenta) ensures efficient anterograde IFT. Trypanosome IFT proteins are concentrated on top of the transition fibres [65] and so KIN2B could be responsible for this concentration. Alternatively, KIN2B could be required for the entry of assembling IFT trains since these have been found at the base of the flagellum but partially on the cytosolic side [61], as also observed in *Chlamydomonas* [9]. At this stage, it is not clear if KIN2B progressively hands over IFT trains to KIN2A, in a situation equivalent to the transition that takes place in the intermediate portion of cilia between the heterotrimeric kinesin-2 and OSM-3 in *C. elegans* [21], or whether IFT trains are released once the transition zone is crossed and then associate again with any of the two kinesins. In the first situation, single molecule imaging revealed that the heterotrimeric kinesin is responsible for progression of IFT trains through the transition zone before being progressively replaced by the homodimeric kinesin OSM-3 in the proximal segment of the axoneme, OSM3 ensuring transport to the tip of the cilium [21]. A similar case could be considered here, but in a shorter portion of the axoneme and between two homodimeric kinesins. In the second situation, trains might “hang around” after they have crossed the transition zone and then be picked up by KIN2A for efficient transport. In both scenarios, we propose that KIN2A has a higher affinity for mature IFT particles, hence once it is associated with them, it runs to the end of the flagellum (**Fig. 9**, large plain magenta arrow) and ensures most anterograde transport of IFT material (0.9 train per second). Nevertheless, a minority of KIN2B molecules remain associated with IFT material (0.4 train per second) and can also transit till the distal end of the flagellum (**Fig. 9**, thin plain blue arrow). The total matches the frequency of IFT trains estimated from monitoring IFT81 anterograde trafficking (1.3 train per second). The remaining KIN2B molecules process along the proximal part of the flagellum but display slower motility, possibly due to negative regulation in the absence of cargo, and detaches from microtubules prematurely without reaching the flagellum tip (**Fig. 9**, dotted blue arrow) before being caught by retrograde trains for recycling. Co-expression of fluorescent KIN2B and IFT140 supports this proposal since KIN2B fluorescent particles that are not associated with IFT140 show slower motion and are limited to the proximal part of the flagellum. However, formal evidence that KIN2A transports IFT particles could not be obtained since co-expression of KIN2A and an IFT protein with two different fluorescent reporters turned out to be impossible. One therefore cannot rule out the possible contribution of other kinesins as observed in *Tetrahymena* [14, 82]. Nevertheless, a third kinesin able to transport IFT particles was not identified, neither by genome mining ([36, 83], this study), nor during the TrypTag project [49]. Therefore, KIN2A and KIN2B must be responsible for all the transport of IFT proteins (in the anterograde direction).

**Figure 9:**
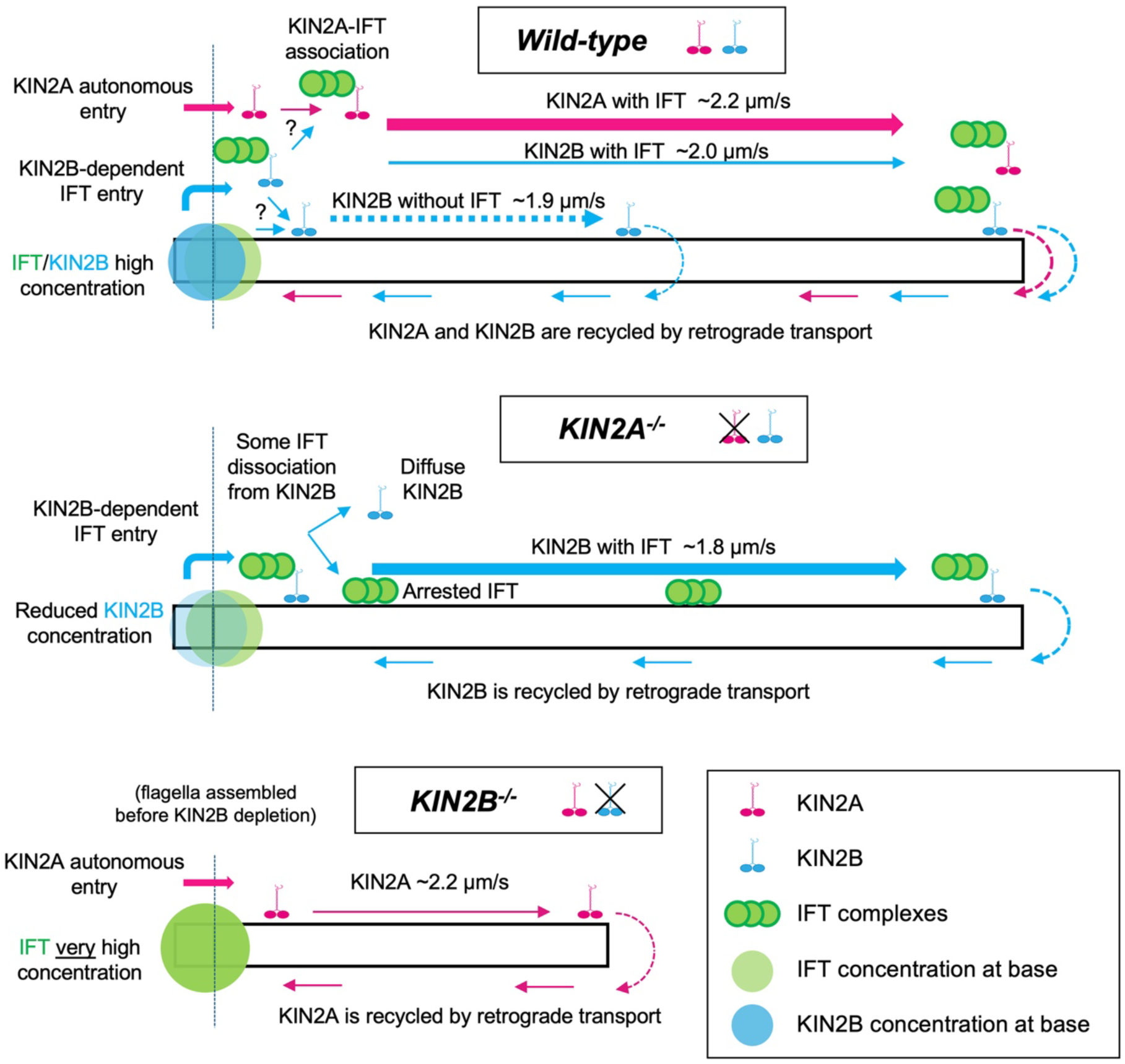
Division-of-labour model for KIN2A and KIN2B. In the wild-type context, we propose that both KIN2A and KIN2B transport IFT trains (plain arrows) with KIN2A ensuring two thirds of the transport (represented by the thicker arrow) while a large proportion of KIN2B travels independently of IFT (shown with the dotted arrow). In the absence of KIN2A, KIN2B is the only motor responsible for IFT transport, but it appears less processive, resulting in frequent arrest of IFT trains. In the absence of KIN2B, flagella are not assembled but those constructed at the previous generation could be examined, revealing that IFT proteins (green) remain stuck at the base while KIN2A can still circulate. This suggests that KIN2B is responsible for IFT train access in the flagellar compartment while KIN2A can enter autonomously. KIN2B at the base is represented by a blue halo whose intensity reflects its relative concentration. The dotted line represents the separation between the cell body and the flagellar compartment. Since the precise localisation of KIN2B at the base could not be demonstrated, it is represented by a circle around the area.

This model can explain the phenotypes following the deletion of each kinesin. In the KIN2A knockout situation (**Fig. 9**, central panel), KIN2B would still be able to transport IFT proteins into the flagellum and since there is no longer competition with KIN2A, it could now ensure their delivery throughout the flagellum, albeit at a slower motion. However, the proposed lower affinity for IFT association would explain the more chaotic transport of the reporter IFT81, with frequent pauses in trafficking. This alteration results in the formation of a shorter flagellum, as expected upon reduction of IFT trafficking [33, 71, 84]. By contrast, the model would predict that in the absence of KIN2B (**Fig. 9B**, bottom panel), IFT proteins cannot enter the flagellum and hence flagellum assembly should be inhibited as observed upon depletion of IFT proteins [69, 71]. The impact of KIN2B depletion could be monitored in flagella assembled in the previous generation, revealing that KIN2A can access the flagellar compartment in the absence of KIN2B (**Fig. 9**). However, IFT proteins are not incorporated and hence pile up at the flagellar base.

KIN2A and KIN2B appear to function the same way in *T. brucei* and *L. mexicana*, suggesting that the division-of-labour system was already in place at least in their last common ancestor and is likely used across trypanosomatids. The major difference is the length of the proximal portion where high amounts of KIN2B are detected, which is much longer in *T. brucei* (compare **Fig. 5** and **Fig. S4**). This could reflect differences in the KIN2B properties or in the anatomy of the flagellum along its length between the two species. There is existing support for the later: outer dynein arms are anchored to microtubules by two different docking complexes whose composition varies along the length of the flagellum. The proximal docking complex is much shorter in *L. mexicana* [59], in proportions quite similar to those observed for KIN2B here. This does not mean that the docking complex regulates KIN2B interactions with microtubules, but it might be a marker of a region of the axoneme with different molecular properties.

Using a combination of FIB-SEM and expansion microscopy, it was recently shown that IFT trains present at the very proximal part of the flagellum (first 1-1.5 µm) look thinner and contain less IFT material than “mature” IFT trains encountered along the rest of the flagellum. Moreover, these trains can be found on all doublet microtubules, while mature trains are concentrated on doublets 3-4 and 7-8 [65]. Possibly KIN2B accesses all doublet microtubules while KIN2A is limited to doublets 3-4 and 7-8. Since transport of IFT trains mostly happens on these specific doublets, KIN2A would therefore be the dominant motor. Indeed, a simple calculation shows that from a total of 0.9 KIN2A and 1.2 KIN2B particles per second, 0.225 KIN2A per second are predicted per doublet (0.9 divided by 4 if KIN2A uses exclusively these 4), while only 0.133 KIN2B per second would be found (1.2 divided by 9, assuming that KIN2B uses equally the 9 doublets equally). So KIN2B would have the potential to carry twice less mature IFT trains on doublets 3-4 and 7-8, a value close to what was observed when comparing with KIN2B particles associated to IFT140 (Fig. 6). This would imply that KIN2B carrying IFT material present on the other doublets would transfer it to KIN2A and continue without cargo before detaching from microtubules and being recovered by retrograde trains. Formal evidence for this model requires high-resolution of KIN2A and KIN2B positioning on microtubule doublets in the proximal portion of the flagellum. Unfortunately, the signal for mNG fusion proteins or the staining with the anti-KIN2B antibodies was too diffuse to obtain conclusive data using expansion microscopy or super-resolution microscopy.

This division-of-labour model can be related to the situation encountered in *C. elegans*, where the homodimeric kinesin OSM3 exhibits fast IFT and is present mostly within cilia, while its heterotrimeric counterpart is concentrated at the transition zone, where it would ensure the passage of IFT through that dense region, and the axoneme proximal portion [21]. KIN2B could therefore be more akin to the heterotrimeric kinesin- 2, while KIN2A would be reminiscent of metazoan homodimeric kinesins such as OSM3 and KIF17. Trypanosomatids belong to the supergroup Discoba but were previously considered ’excavates’, which proved to be a paraphyletic grouping of organisms [85]. Either group was often considered close to the root of the eukaryotic tree, therefore possessing many features of the last eukaryotic common ancestor (LECA). Indeed, a recent study proposed that the LECA was of ’excavate’-like nature [86]. In this context, one could propose that heterotrimeric kinesins emerged following the duplication of a KIN2B-like ancestor and the acquisition of the KAP. This would have allowed gathering the functions of IFT entry and trafficking, rendering homodimeric kinesins less critical for ciliogenesis, which could potentially explain their loss as observed in algae. Nevertheless, liberating a homodimeric kinesin from its role in IFT could have allowed the emergence of novel functions [87], such as the transport of signalling molecules, as reported for KIF17 in photoreceptors or olfactory neurons [88, 89]. In *T. brucei*, the proximal KIN2B particles not associated with IFT proteins display slower motion than those carrying IFT cargoes. However, their speed is equivalent to that of KIN2B molecules whose candidate regulatory tail was replaced with only the GCN4 dimerization domain and so faster than the partially inhibited motor. This suggests that these particles could transport distinct cargoes whose identity and function remain to be discovered. This could be relevant in the context of signalling, where the flagellum has been proposed to be a critical actor [90–92]. A different option would consider that the combination of heterotrimeric (with KAP) and homodimeric kinesin was already present in the LECA, but that heterotrimeric kinesin, including KAP, was lost in the common euglenozoan ancestor, leading to a redistribution of its roles between KIN2A and KIN2B.

## Conclusion

In summary, our multi-disciplinary work reveals for the first time that anterograde IFT and flagellum formation can be carried out solely by homodimeric kinesins, which work cooperatively to ensure entry of the large IFT complexes into the flagellar compartment and their circulation afterwards. In the future, it will be exciting to untangle how these motors associate with IFT trains and with microtubule doublets, especially in the peculiar environment of the trypanosome flagellum where IFT is initiated on all nine doublets before being restricted to only four of them [61, 65].

## Materials and Methods

### Homology searches and phylogenetic analysis

Homology searches were performed by blast+ v2.9.0 [93] integrated in the AMOEBAE workflow [94] using sequencing data from NCBI (https://www.ncbi.nlm.nih.gov/), TriTrypDB [95], EukProt [96], JGI (https://jgi.doe.gov/), UniProt (https://www.uniprot.org/), and previous publications [97–100]. Identified kinesin-2 sequences were aligned together with reference kinesin sequences from a previous study [36] using MAFFT v7.520 [101] under the L-INS-i strategy. Poorly aligned positions were removed by trimAl v1.4.rev15 [102] using -gt 0.8 option. Maximum-likelihood phylogenetic tree was performed in IQ-TREE v3.0.1 [103] with the LG+G model for the guide tree, followed by PMSF analysis [104] using the LG+C50+G4 model and the guide tree input with 1,000 replicates for ultrafast bootstraps [105] and a maximum of 10,000 iterations.

### DNA constructs, design and virus generation for *in vitro* expression

All the DNA sequences used in the study were obtained from published information in the following data bases: NCBI, TrypTag (http://www.tryptag.org/)[95] and LeishGEdit (https://www.leishgedit.net) [106], and are listed in **Table S2**. They were synthesised (GenScript Biotech) and cloned into the pFastBac-1 vector according to the manufactureŕs instructions (Thermo Fisher Scientific). The constructs were either FLAG- or 6xHis-tagged to facilitate protein purification. Recombinant bacmid generation and virus amplification were done according to the manufactureŕs instructions (Thermo Fisher Scientific).

### Protein expression, purification, and fluorescent labelling

All the proteins were expressed using the Baculovirus Expression system in Sf9 insect cells (*Spodoptera frugiperda*) according to the manufactureŕs instructions (Thermo Fisher Scientific). The proteins were expressed in suspension cultures with a concentration of 2x10^6^ cells/ml cultured in Sf-900^TM^ II SFM medium supplemented with 10% (v/v) foetal bovine serum and 0.5% (w/v) gentamicin. The cultures were infected with 2-8% (v/v) appropriate virus suspension and incubated for approximately 60 hours at 28°C and 110 rpm. After incubation, cells were harvested by centrifugation at 2,500 *g* for 15 minutes.

The expressed proteins contained either N- or C-terminal FLAG (DYKDDDDK) or 6xHis-tags and were purified through FLAG-tag or His-tag affinity purification. In FLAG-tag affinity purification the cells were lysed with 4% (v/v) of FLAG lysis buffer (50 mM PIPES pH 6.9, 300 mM potassium acetate, 1 mM MgCl_2_, 1 mM DTT, 0.1 mM ATP, 0.5% Triton X-100, 10% glycerol) and Complete Protease Inhibitor Cocktail (Roche), and centrifuged at 65,000 *g* and 4°C for 10 min. The supernatant containing the protein of interest was then incubated with 4% (v/v) of anti-FLAG MS Affinity Agarose gel (Sigma-Aldrich) for 90 minutes at 4°C tumbling end-over-end. The FLAG beads were washed three times with FLAG wash buffer I (80 mM PIPES pH 6.9, 500 mM potassium acetate, 1 mM MgCl_2_, 1 mM DTT, 0.1 mM ATP, 0.1% Tween-20, 1 mM EGTA, 10% glycerol) and three times with FLAG wash buffer II (80 mM PIPES pH 6.9, 200 mM potassium acetate, 1 mM MgCl_2_, 1 mM DTT, 0.1 mM ATP, 0.1% Tween-20, 1 mM EGTA, 10% glycerol). For fluorescent labelling, the respective tagged proteins were incubated with 10 µM of either HaloTag^®^ Alexa Fluor^®^ 488/Alexa Fluor^®^ 660 (Promega) or SNAP surface^®^ Alexa Fluor^®^ 488/Alexa Fluor^®^ 647 (New England Biolabs) dyes in FLAG wash buffer II for 45 minutes at 4°C tumbling end-over-end and protected from the light. The beads were then extensively washed with FLAG wash buffer II and eluted in 1-2 times (v) of the resin in FLAG elution buffer (FLAG wash buffer II containing 0.5 mg/ml FLAG peptides; Sigma-Aldrich).

In His-tag purification, cells were lysed in 4% (v/v) His-lysis buffer (50 mM PIPES pH 8.0, 300 mM potassium acetate, 10 mM imidazole, 0.5% Triton-X 100, Complete Protease Inhibitor Cocktail (Roche), 1 mM DTT, 10% glycerol). The lysate was centrifuged as in FLAG-tag purification and the supernatant was incubated with 4% (v/v) of pre-washed Ni-NTA coated Sepharose beads (Qiagen) for 90 minutes at 4°C tumbling end-over-end. After incubation, the beads were washed four times with His wash buffer (50 mM PIPES pH 8.0, 500 mM potassium acetate, 40 mM imidazole, Complete Protease Inhibitor Cocktail (Roche), 1 mM DTT, 10% glycerol) and were eluted in 1-2 times (v) of the resin in His elution buffer (50 mM PIPES pH 7.5, 100 mM potassium acetate, 500 mM imidazole, 1 mM EGTA, Complete Protease Inhibitor Cocktail (Roche), 1 mM DTT, 10% glycerol).

### Single-molecule transport assays

For transport assays, motor proteins were fluorescently labelled and purified through a FLAG-tag affinity purification. The biotinylated and fluorescently labelled microtubule filaments were initially attached to the glass surface of a flow chamber using biotin-streptavidin interaction as described previously [40]. The motor proteins were then diluted in the motility buffer (10 mM PIPES pH 6.9, 2 mM MgCl_2_, 1 mM EGTA, 5 mM DTT, 2 mM ATP, 0.18 mg/ml glucose-oxidase (G2133; Sigma-Aldrich), 0.06 mg/ml catalase (C3155; Sigma-Aldrich), 0.4% glucose) and flushed into the flow chamber. The proteins were excited by laser beams with the appropriate wavelength and tracked using the objective-type Leica DMi8 TIRF microscope (Leica) equipped with a Plan Apochromat 100×/1.47 NA oil objective (Zeiss) and an Andor iXon Ultra EMCCD camera (Andor Technology). The run length and velocity data were analysed using a custom-written MATLAB (MathWorks Inc.) routine.

### *In vitro* co-localisation assays

The *in vitro* co-localisation assays were conducted as described previously [80]. Briefly, the purified proteins were fluorescently labelled with appropriate dyes according to their fluorescent tags as described above. After purification, they were mixed in equimolar concentration and were incubated in a rolling incubator overnight at 4°C. The protein mixtures were appropriately diluted and pipetted directly onto the glass slide and covered with a cover slip. Images were acquired using the TIRF microscope as explained previously. Co-localisation efficiency of the labelled proteins was analysed using a custom-written MATLAB (MathWorks Inc.) routine.

### Size-Exclusion Chromatography coupled to Multiple-Angle Light Scattering (SEC-MALS)

SEC-MALS analysis was conducted to determine the oligomeric state of the *T. brucei* motor proteins. The motor proteins were analysed using a Superose 6 Increase 10/300 GL column (GE Healthcare) with a flow rate of 0.5 ml/min, controlled by an injector pump (Agilent 1260 Infinity series). The column was initially equilibrated with two volumes of gel filtration buffer (25 mM PIPES pH 7.0, 200 mM NaCl, 1 mM MgCl_2_, 1 mM EGTA, 1 mM DTT). Protein samples were manually loaded into the sample loop and injected into the column at the start of the run. The elution was analysed using a UV absorbance detector (Agilent 1260 Infinity series) set at 280 nm wavelength and by a MALS detector (DAWN8; Wyatt Technology) which was connected inflow to the column. The final eluent was collected in separate fractions using a fraction collector (Agilent 1260 Infinity series), and the fractions were subsequently analysed using SDS PAGE. The molar mass of motor proteins was determined using the ASTRA v8.2.2 software (Wyatt Technology).

### Cell culture

Procyclic forms of *T. brucei* strain 427 and TREU927 containing the plasmid pJ1339 [107] were grown in SDM79 medium supplemented with 10% FCS [108] at 28°C. Culture density was maintained between 5x10^5^ and 10^7^ cells/mL. Promastigote forms of *L. mexicana* derived from WHO strain MNYC/BZ/62/M379, expressing Cas9 and T7 RNA polymerase were grown in M199 (Life Technologies) supplemented with 2.2 g/L NaHCO_3_ (Sigma-Aldrich, S6014), 0.005% hemin (Sigma-Aldrich, H9039), 40 mM HEPES HCl pH 7.4 (Merk Life Science, H3375) and 10% FCS (Merk Life Science, F9665) at 28 °C. Culture density was maintained between 10^5^ and 10^7^ cells/mL for continued exponential population growth.

### Generation of endogenous tagging constructs for *in vivo* expression and gene deletion constructs

Constructs for *T. brucei* IFT81 (Tb927.10.2640), KIN2A (Tb927.5.2090) and KIN2B (Tb927.11.13920) endogenous tagging with mNeonGreen (mNG) at the N- or C-terminus were done as described [44]. For the KIN2A and KIN2B cell lines with both alleles tagged, the same primers were used to generate constructs containing puromycin or hygromycin resistance genes, followed by allele tagging under selection with the respective antibiotic. For the double-tagged cell line, cells expressing mNG::KIN2B were nucleofected with the plasmid p2845TdTomatoIFT140 [54] linearised with MfeI, allowing the expression of IFT140 (Tb927.10.14470) fused to tdTomato (tdT) at the N-terminus from its endogenous locus. Constructs for *KIN2A* gene deletion mediated by CRISPR-Cas9 were generated as described [58].

For *L. mexicana*, endogenously tagged and deletion mutant cell lines were generated by transfection of Cas9T7 with the appropriate combination of DNA encoding sgRNAs and repair constructs with flanking 30-base homology arms carrying the necessary drug-selectable marker and, for tagging, the fluorescent protein, as previously described [58]. All the primers used for the construct generation are listed in the **Table S2**.

### Electroporation and drug selection

Prior to electroporation, the necessary sgRNA and repair construct PCR reactions were mixed, then precipitated with 10 µL of 3 M sodium acetate (Sigma, S8750) and 300 µL of EtOH at -80 °C for 20 minutes, pelleted by centrifugation (20,000 *g*, 30 minutes at 4 °C), washed once with 70% EtOH, re-pelleted, and then resuspended in 10 µL of water. *L. mexicana* was electroporated using the Nucleofector II (Amaxa) in 2 mm cuvettes with the X-100 program. 1x10^7^ cells were mixed with 250 µL electroporation buffer (a ratio of 1:6:3 of 1.5 mM CaCl_2_ (Merck, 202940) and 200 mM Na_2_HPO_4_ (Merck, 71640), 70 mM NaH_2_PO_4_ (Merck, S5011), 15 mM KCl (Merck, P5405), 150 mM HEPES pH 7.4 (Merck, H4034). Cells were immediately resuspended in pre-warmed M199. Successful transfectants were selected and maintained using the necessary combination of drugs selection (20 μg/mL puromycin dihydrochloride (Puro; Cambridge Science Ltd), 5 μg/mL blasticidin S hydrochloride (Bla; Stratech, B4879) and 40 μg/mL G418 disulfate (Neo; Invitrogen). Drugs were added 15 hours after transfection, and resistant populations took 4 to 15 days to emerge.

### Protein expression and antibody generation

A *KIN2B* 664bp gene fragment (Tb927.11.13920, bp :1467-2129) flanked by BamHI (5’) and HindIII (3’) sites was synthesised (Genecust) and cloned into the corresponding restriction sites of the pGEXB expression plasmid. pGEXB-Kin2B plasmid was sequenced to confirm identity and correct fusion with Glutathion-S-Transferase (GST). pGEXB-KIN2B plasmid was transformed into the competent BL21 strain of *E. coli*. Following IPTG induction, *E. coli* cells were harvested by centrifugation, resuspended in BugBuster lysis reagent (Novagen) supplemented with lysozyme (1kU /ml) at room temperature. Protein expression was analysed by SDS-polyacrylamide gel electrophoresis followed by Coomassie staining. As most of the recombinant KIN2B GST protein was insoluble, additional solubilisation step in 2M urea was performed prior purification on 2M urea equilibrated glutathione agarose as described [109]. 20µg was injected into BALB/C mice for immunisation. Mice were housed in the Institut Pasteur animal facilities, which are accredited by the French Ministry of Agriculture for performing experiments on live mice, in compliance with the French and European regulations on care and protection of laboratory animals (European Directive 86/609/UE). Protocols for immunisation were approved by the veterinary staff of the Institut Pasteur animal facility and were performed in compliance with National Institutes of Health Animal Welfare Assurance A5476-01 issued on 7 February 2007.

### Diagnostic PCR and whole-genome sequencing for validation of gene deletion in *T. brucei*

For the production of the *KIN2A^-/-^* cell line, the loss of the target open reading frame (ORF) in drug-resistant transfectants was verified by a diagnostic PCR using primers designed to amplify a 1000 bp fragment within the target gene’s ORF. Genomic DNA from the parental cell line was used as a positive control for the amplification of the target ORF. To confirm the correct integration of the drug-resistance cassettes, specific primers were used to target the 5’UTR of the target ORF in combination with either the *hygR* or *neoR* genes. Each PCR reaction consisted of 100 ng of genomic DNA, 1 μL each of 10 μM forward and reverse primers, and 25 μL of GoTaq® Master Mix (Promega, M7123) in a final reaction volume of 50 μL. The thermocycling profile consisted of an initial denaturation at 94°C for 5 minutes, followed by 30 cycles of denaturation at 94°C for 30 seconds, annealing at 60°C for 30 seconds, and extension at 72°C for 70 seconds, with a final elongation step at 72°C for 10 minutes.

Genomic DNA from the parental and drug-resistant *T. brucei* cell lines was extracted using the DNeasy Blood & Tissue Kit (Qiagen, 69506). Library preparation and paired-end sequencing were performed by BGI Genomics (China) with a target average depth of 40x. Raw sequencing reads were processed using fastp [110] to remove adapter sequences, low-quality bases (Phred score < 20), and short reads. Cleaned reads were aligned to the *T. brucei* TREU927 reference genome (v7.0, TriTrypDB) using BWA-MEM. To confirm the successful knockout of the target ORF, alignment files (BAM) were visualised using the Integrative Genomics Viewer (IGV) to verify the absence of reads at the target locus. To assess off-target effects and secondary mutations associated with drug resistance, *de novo* variants (SNPs and Indels) in the drug-resistant cell line were identified by filtering out all baseline variants present in the parental strain.

### Diagnostic PCR for validation of gene deletion for *L. mexicana*

To verify the loss of the target open reading frame (ORF) in drug-resistant transfectants, a diagnostic PCR was performed using primers that amplify a short fragment (100–300 bp) within the target gene’s ORF. Genomic DNA from the parental cell line served as a positive control to confirm successful amplification of the target ORF. If no PCR product was detected, the annealing temperature was reduced by up to 2°C to optimise primer binding. As a technical control, a short fragment of the *PF16* ORF (LmxM.20.1400) was amplified to confirm the presence of genomic DNA in the knockout cell line. Each PCR reaction contained 25–100 ng of genomic DNA, 1.25 μL each of 10 μM forward and reverse primers, and 12.5 μL of FastGene Optima HotStart Ready Mix with dye (Nippon Genetics, P8-0082), with water added to a final volume of 25 μL. The thermocycling conditions were: initial denaturation at 95 °C for 3 minutes; 35 cycles of 95 °C for 15 seconds; 58 °C for 15 seconds; and 72 °C for 30 seconds; followed by a final elongation at 72 °C for 1 minute.

### Cas9-marker-free gene disruption

For marker-less gene disruption of KIN2B and IFT172, a Custom Alt-R™ CRISPR- Cas9 toolkit was used. Guide RNA and repair template design were adapted from [72].

To disrupt gene expression, a guide RNA in the first 400 bp of the ORF was chosen that had the least homology with the rest of the *T. brucei* genome. Then, a repair cassette was designed that contained six stop codons in the centre. To target all ORFs, thymine nucleotides were inserted between stop codons to interrupt each reading frame. For diagnostic purposes, a BamHI restriction site was introduced in between the two series of stop codons. Both sides of the stop codon insert were flanked by 50 bp-long homology regions targeting the gene of interest. These homology arms were spaced 23 nucleotides apart and flank the region that was targeted by the guide RNA. In a successfully edited genomic sequence, the targeting region of the guide RNA including the PAM sequence was therefore replaced by the stop codon repeat. To generate duplex RNA, an equimolar mix of tracrRNA and crRNA was heated at 94°C for 5 minutes and then left to cool at room temperature. RNA duplex (∼0.12 nmol/reaction) and Cas9 enzyme (30µg/reaction) were mixed and incubated to associate the Cas9 ribonucleoprotein. Each Cas9-RNA-complex was then mixed with 8µg repair template (amplified by PCR followed by phenol-chloroform purification) and transfected with an Amaxa nucleofector II from Lonza (program X-014). Alt-R™ CRISPR-Cas9 crRNA, Alt-R™ CRISPR-Cas9 crRNA, Alt-R™ *Streptococcus pyogenes* Cas9 Nuclease V3, as well as the repair template for PCR amplification (MiniGenes in pUCIDT (Amp) Golden Gate vectors) were ordered from Integrated DNA Technologies. The nucleofected cells were observed under the microscope after 4 to 48 hours by live fluorescence microscopy as well as after IFA with antibodies recognising either the Ty-1-tag (BB2)[45], IFT172 [52], the axonemal marker TbSAXO (Mab25)[51] or the FTZC transition zone marker [50].

### Western blot

The lysate equivalent to 1×10^6^ cells was loaded onto a 5-15% gradient SDS–PAGE gel. After transfer, membranes were incubated with blocking solution (3% milk, 0.05% Tween 20 in PBS) for 1 h, followed by incubation with neat BB2 (anti-Ty tag) or anti-ALBA 1:1000 [47] in blocking solution for 1 hour under agitation. After three washes in 0.05% Tween 20 in PBS, membranes were incubated with secondary mouse IgG HRP-linked antibody (Amersham NA931) diluted 1:10000 in blocking solution for 1 h. Proteins were detected by ECL using the ImageQuant 800 system (Amersham).

### Immunofluorescence microscopy

For *T. brucei*, cells were harvested, resuspended in TDB (5 mM KCl, 80 mM NaCl, 1 mM MgSO_4_, 20 mM Na_2_HPO_4_, 2 mM NaH_2_PO_4_, 20 mM glucose, pH 7.4) and settled onto poly-L-lysine-covered glass slides for 10 minutes. Cells were fixed in 4% formaldehyde (Electron Microscopy Sciences) in PBS for 5 minutes, followed by a 5- minute incubation in -20°C methanol. After rehydration for 10 minutes in PBS, samples were incubated for 1 hour in primary antibodies diluted in blocking solution (1% BSA in PBS) for 1 hour. The following primary antibodies were used: BB2 [45] diluted 1:10, anti-IFT172 [52] diluted 1:50, mAb25 [51] diluted 1:10 and anti-FTZC [50] diluted 1:500. Cells were washed three times in PBS, then incubated in secondary antibodies diluted in blocking solution, and washed three times in PBS. After 5-minute incubation in 20 μg/ml Hoechst 33342 (Sigma-Aldrich) or DAPI in PBS for DNA staining, cells were washed in PBS and mounted in ProLong™ Gold (ThermoFisher Scientific) before imaging.

For *L. mexicana*, cells expressing fluorescent fusion proteins were imaged live. Cells were washed three times by centrifugation at 800 *g*, followed by resuspension in PBS. DNA was stained by including 10 μg/mL Hoechst 33342 in the second washing. Washed cells were settled on glass slides and were observed immediately. Widefield epifluorescence and phase-contrast images were captured using a Zeiss Axioimager.Z2 microscope with a 63×/1.40 numerical aperture (NA) oil immersion objective and a Hamamatsu ORCA-Flash4.0 camera.

### Live cell imaging and IFT analysis

For *T. brucei*, live cells were washed and kept in DMEM fluorobrite medium for live imaging. Before the acquisition, 3 µL of concentrated cells were spread on a glass slide and covered with a 1.5H coverslip. The samples were kept in a temperature-controlled chamber at 27°C. Time-lapses were taken with a spinning-disk confocal microscope (UltraView VOX; PerkinElmer) equipped with an EMCCD camera (C-9100; Hamamatsu Photonics) operating in streaming mode and using a Plan Apochromat 100x/1.57 NA oil objective (ZEISS). Sequences of 15 or 30 seconds were acquired with an exposure time of 100 ms per frame at maximum acquisition speed, using Volocity software. Kymograph analysis was done using Icy software v2.5 as described [39, 111].

After kymograph extraction, the Icy software was used for the image background subtraction of green and red anterograde kymographs, followed by the generation of the Manders’ coefficients between the two signals with the Colocalization Studio plugin, using the same regions of interest as for the speed and frequency measurements.

Epifluorescence of *L. mexicana* cell lines were carried out using cells from late exponential growth phase (5 × 10^6^ to 1 × 10^7^ cells/ml). A 500 μl sample of culture was washed by centrifugation at 1200 g and resuspended three times in PBS, then resuspended in 50 μl PBS and 1 μl was placed on a plain glass slide with a coverslip. Videos were captured at room temperature using an Axio Observer A1 microscope (Zeiss) using a 63× 1.40 NA objective at 5 frames per second. Flagella were traced from maximum signal intensity projection of the mNG signal from all frames of the video. For background subtraction, a fast Fourier transform filter with a maximum structure size of 5 px followed by a rolling ball background subtraction with a radius of 10 px was used, and kymographs were generated from the background-subtracted image. Signal intensity along the flagellum from the basal body to the flagellum tip was measured from the average of the first 10 frames of the raw (non-background subtracted) video. The line was traced from the basal body kinesin signal when visible, or 2.5 μm into the anterior end of the cell, the nominal depth of the flagellar pocket, when not visible. *Leishmania* flagella have variable length, so signal intensity was measured in 20 even bins along the flagellum length and normalised to the maximum signal intensity for that cell.

### Transmission Electron Microscopy

For *T. brucei*, procyclic cells were fixed overnight at 4°C in a solution containing 2.5% glutaraldehyde and 2% formaldehyde in 0.1 M PHEM (PIPES 12 mM, HEPES 5 mM, EGTA 2mM, MgSO_4_ 0.8 mM) buffer. After fixation, cells were washed three times with 0.1 M PHEM buffer 5 min each and post-fixed with 1% osmium and 1.5% potassium ferrocyanide in 0.1 M PHEM buffer for 1 hour. Samples were treated for 30 min with 1% tannic acid and 1 h with 1% osmium tetroxide, rinsed in water and incubated in 1% uranyl acetate dissolved in 25% ethanol for 30 min. Cells were dehydrated in an ethanol series comprising 25%, 50%, 75% and 95% ethanol solutions (5 minutes each), followed by three immersions in 100% ethanol (10 minutes each). Cells were embedded in epoxy resin after 48 hours at 60°C of polymerization. Sections of 50-70 nm were obtained using an EM UC6 ultra-microtome (Leica) and stained with 2% uranyl acetate in milli-Q water and 80 mM lead citrate in milli-Q water before imaging. The sample was imaged using a Tecnai BioTWIN 120 microscope (FEI) equipped with a MegaView II camera (Arecont Vision).

For *L. mexicana*, cells were fixed directly in medium for 10 minutes at room temperature in 2.5% glutaraldehyde (glutaraldehyde 25% stock solution, EM grade, Electron Microscopy Sciences). Centrifugation was carried out at room temperature for 5 minutes at 16,000 *g*. The supernatant was discarded and the pellet was fixed in 2.5% glutaraldehyde and 4% PFA (16% stock solution, EM grade, Electron Microscopy Sciences in 0.1 M PIPES pH 7.2) for a minimum of 2 hours. Cells were embedded in 3% agarose and contrasted with 1% OsO_4_ in water (osmium tetroxide 4% aqueous solution, Taab Laboratories Equipment) for 2 hours at 4 °C. Cells were stained with 2% of uranyl acetate for 2 hours 4 °C. After serial dehydration with ethanol solutions, samples were embedded in low-viscosity resin Agar 100 (Agar Scientific, UK) and left to polymerise at 60 °C for 24 hours. Ultrathin sections (90 nm) were collected on nickel grids using a Leica EM UC7 ultramicrotome and stained with 1% uranyl acetate in water (uranyl acetate dihydrate, Electron Microscopy Sciences) and Reynolds lead citrate [73] (80 mM Lead nitrate (Thermo Fisher, L/1450), 120 mM sodium citrate (Sigma, 71405) and 1 M sodium hydroxide (SLS, CHE3422). Observations were made on a Thermo Fisher Scientific Tecnai12 or JEOL 2100 Plus 200 kV transmission electron microscope with a Gatan OneView camera.

## Statistical tests

The statistical test used for Figures 5, 6 and 7 was a two-way ANOVA with Tukey’s multiple comparisons test. For IFT trafficking data, a total of 60 cells were analysed (20 cells per experiment, in triplicate). Train number varied between 512 and 1003, according to the cell line since kinesins and IFT proteins display different frequencies.

## Acknowledgements

We are grateful to the Photonic Bioimaging and to the Ultrastructural Bioimaging facilities of the Institut Pasteur for access to their equipment. We thank Dr. Errin Johnson for technical assistance for transmission electron microscopy (Electron microscopy facility at Sir William Dunn School of Pathology, Oxford University, United Kingdom). The *Leishmania* aspect of this work was initiated in the laboratory of Professor Keith Gull (University of Oxford) and we thank him for his support and ideas.

Work at the Institut Pasteur was supported by La Fondation pour la Recherche Médicale (EQU202203014654), the ANR (ANR-18-CE13-0014-01) and a French Government Investissement d’Avenir programme, Laboratoire d’Excellence “Integrative Biology of Emerging Infectious Diseases” (ANR-10-LABX-62-IBEID). ZÖ and AC were funded by the Deutsche Forschungsgemeinschaft (DFG, German Research Foundation) – OE 501/8-1 and OE 501/5-1. CF and RW were supported by a Wellcome Trust Sir Henry Dale Fellowship [211075/Z/18/Z]. ZW worked in Professor Keith Gull’s laboratory (University of Oxford), which was supported by the Wellcome Trust (104627/Z/14/Z, 108445/Z/15/Z). K.Z. and V.Y were funded by the EU’s Operational Program “Just Transition” LERCO CZ.10.03.01/00/22_003/0000003.

The authors declare that they have no conflict of interest.

---

**Figure S1.**
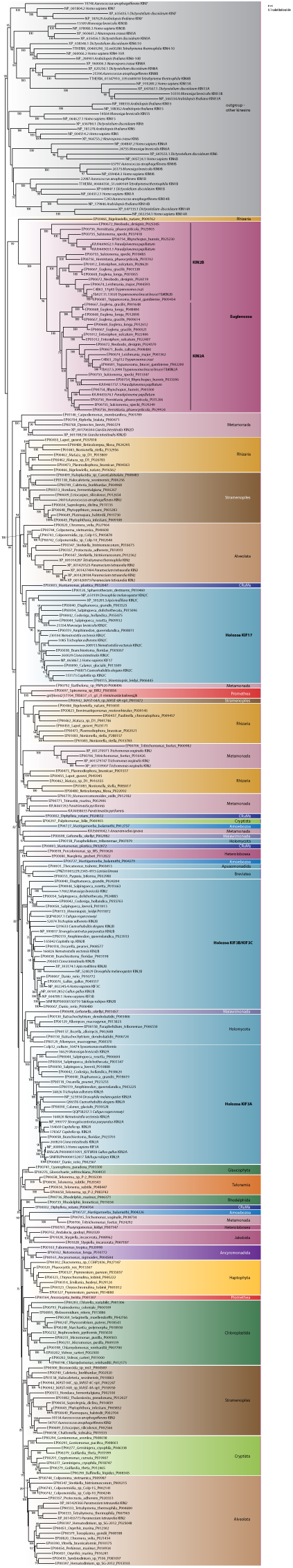
Phylogenetic analysis of kinesin-2. The maximum-likelihood phylogenetic tree was performed in IQ-TREE using PMSF analysis with 1,000 replicates for ultrafast bootstraps (UFB). Only UFB supports ≥75 are shown. Reference kinesin-2 and outgroup kinesin sequences were obtained from [36]. Eukaryotic clades are colour-coded as in Fig. 1.

**Figure S2:**
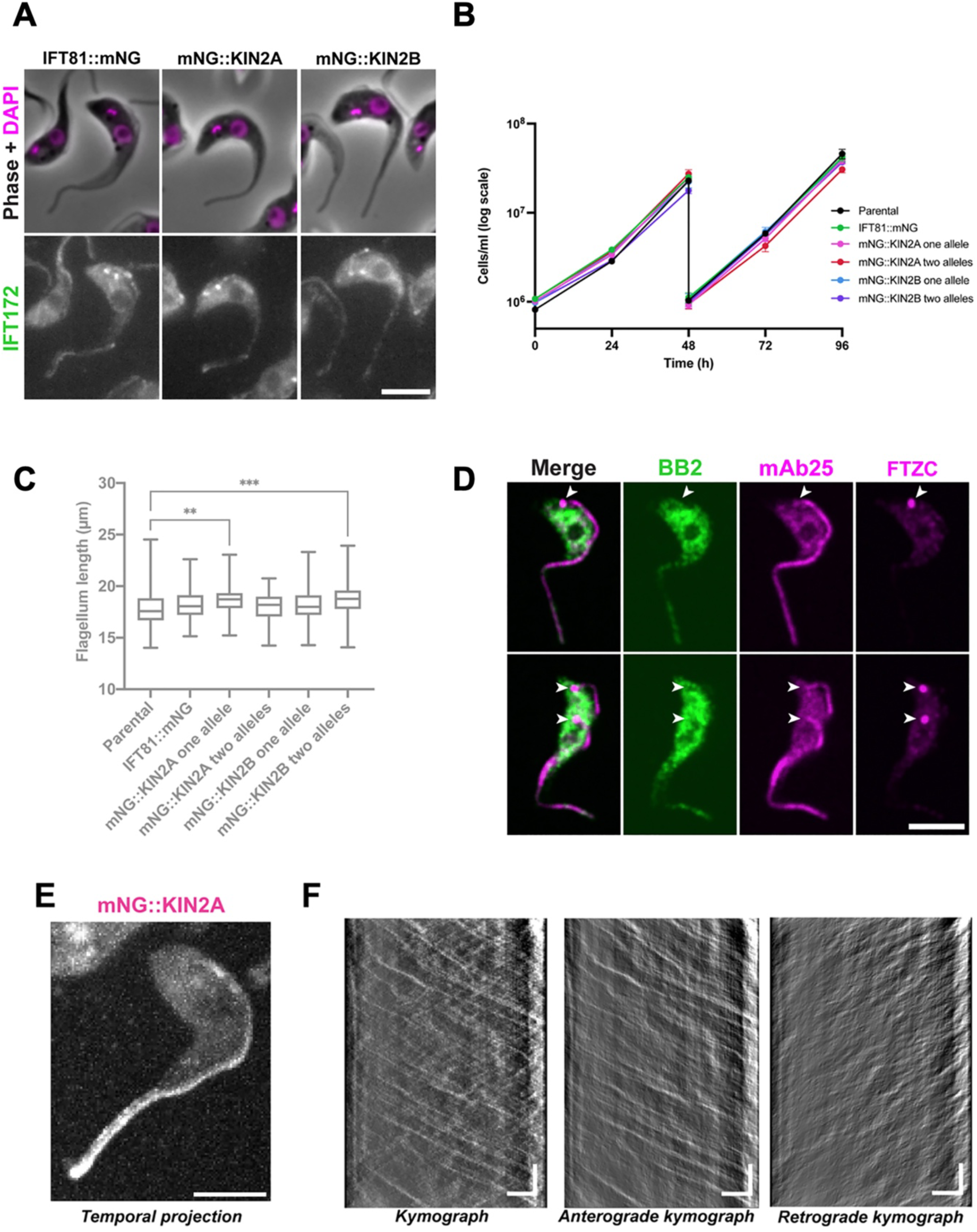
KIN2A and KIN2B tagging in *T. brucei* procyclic form. **(A)** *T. brucei* cells expressing IFT81::mNG, mNG::KIN2A and mNG::KIN2B were stained with anti-IFT172 antibody. Scale bar: 5 µm. **(B)** Growth curves of the cell lines expressing IFT81::mNG, mNG::KIN2A or mNG::KIN2B with single or double alleles tagged. **(C)** Flagellum length measurements of the same tagged cell lines seen in **B**. **(D)** Cells expressing mNG::KIN2A were stained with anti-Ty tag (BB2, green), an antibody against the axonemal protein TbSAXO (mAb25, magenta, third column), and an antibody against the transition zone protein FTZC (magenta, fourth column), showing that KIN2A does not accumulate at the flagellar base (white arrowheads). Scale bar: 5 µm. **(E)** Temporal projection of a cell expressing mNG::KIN2A. Scale bar: 5 µm. **(F)** Kymographs extracted from the cell shown in **E**. Horizontal scale bar: 3 µm; vertical scale bar: 3 s.

**Figure S3.**
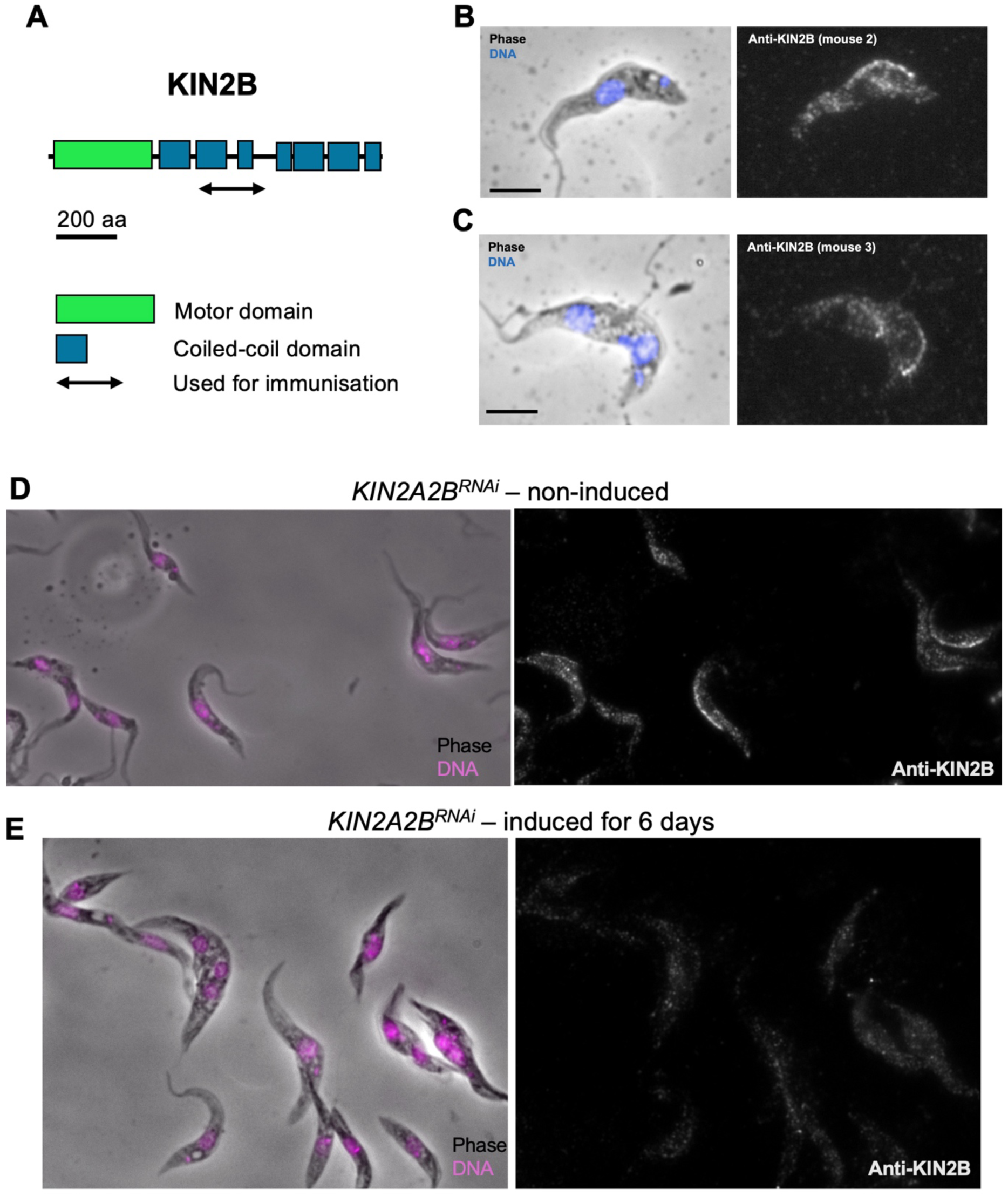
Native KIN2B localises to the flagellar base and to the proximal portion of the flagellum. **(A)** Domain structure of the KIN2B protein and the region used for immunisation. **(B-C)**. Immunofluorescence of wild-type trypanosome stained with the indicated anti-KIN2B mouse antisera. The left panel shows the phase contrast image merged with DAPI (blue) and the right panel shows the anti-KIN2B staining (white). Native KIN2B is concentrated at the base and along the proximal portion of the flagellum, as observed for the mNG::KIN2B fusion protein. **(D-E)** *KIN2A2B^RNAi^* cells were grown without (non-induced) or with (induced for 6 days) tetracycline and stained with an anti-KIN2B. The signal is significantly reduced in RNAi conditions. Please note that a certain level of RNAi leakiness can be observed in this cell line [33], explaining the reduced signal in non-induced conditions compared to wild-type cells.

**Figure S4:**
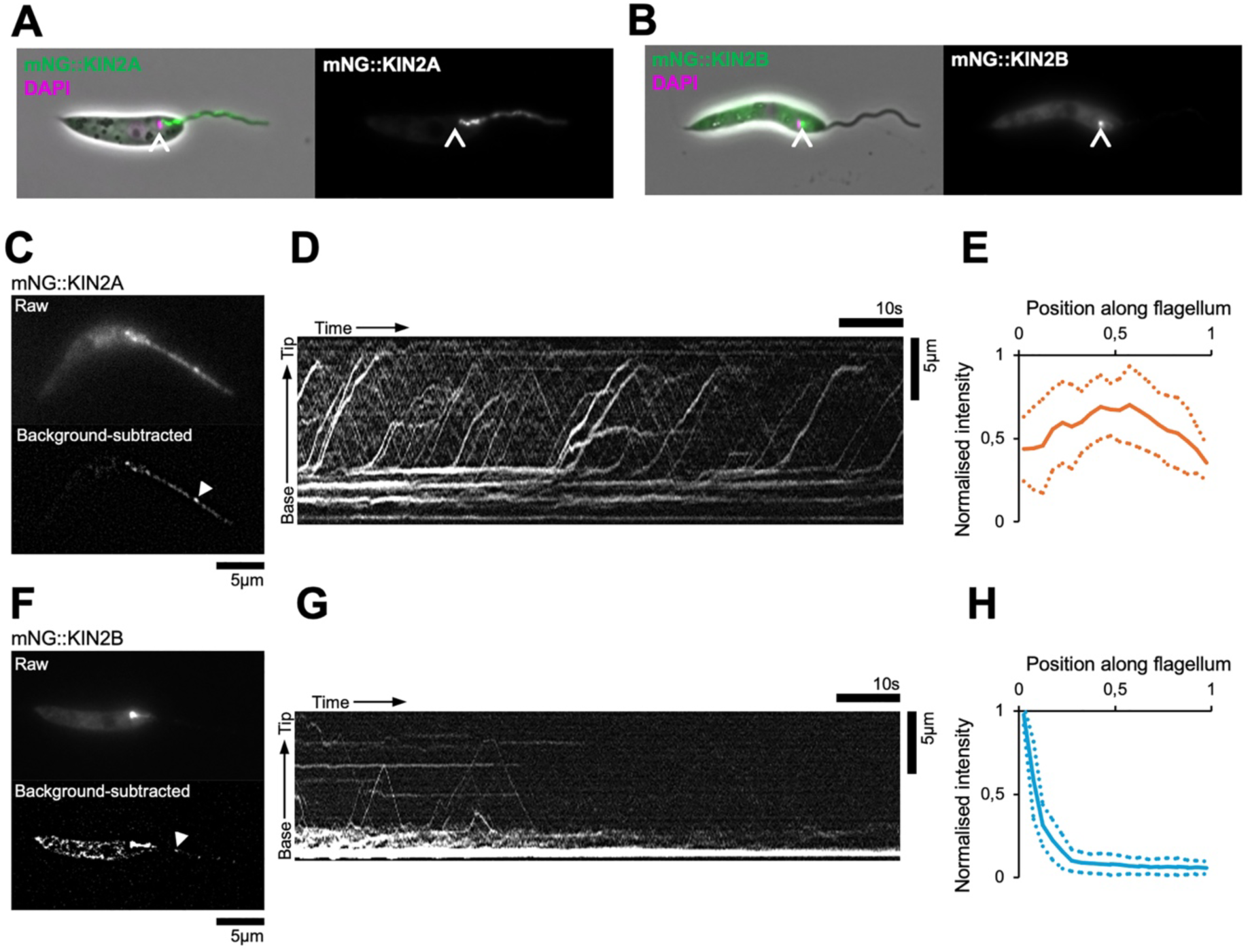
KIN2A and KIN2B in *L. mexicana* share a similar localisation as in *T. brucei.* (A-B) Live epifluorescence images of mNG::KIN2A (A) and mNG::KIN2B (B) in *L. mexicana* promastigotes, stained with DAPI (magenta) showing the location of the fluorescent fusion proteins on merged images (green, left) or alone (white, right panels). The position of the kinetoplast (arrowhead) indicates the base of the flagellum. No particular concentration can be detected for mNG::KIN2A (A), while a clear spot is visible for mNG::KIN2B (B). (C,F) Still epifluorescence images of mNG::KIN2A (**C**) and mNG::KIN2B (**F**) in *L. mexicana* promastigotes. The top panel shows the raw fluorescence micrograph. The bottom panel shows a background-subtracted image to enhance small structures, with an example IFT particle indicated with an arrowhead. **(D,G)** Kymographs of the cells shown in A, showing mNG::KIN2A (top) and mNG::KIN2B (bottom). **(E,H)**. Fluorescent signal distribution along the flagellum, from basal body to flagellum tip, for mNG::KIN2A (**E**) and mNG::KIN2B (**H**). Solid line represents the mean and the dotted line represents the standard deviation of *n* = 12 and 7 cells respectively.

**Figure S5:**
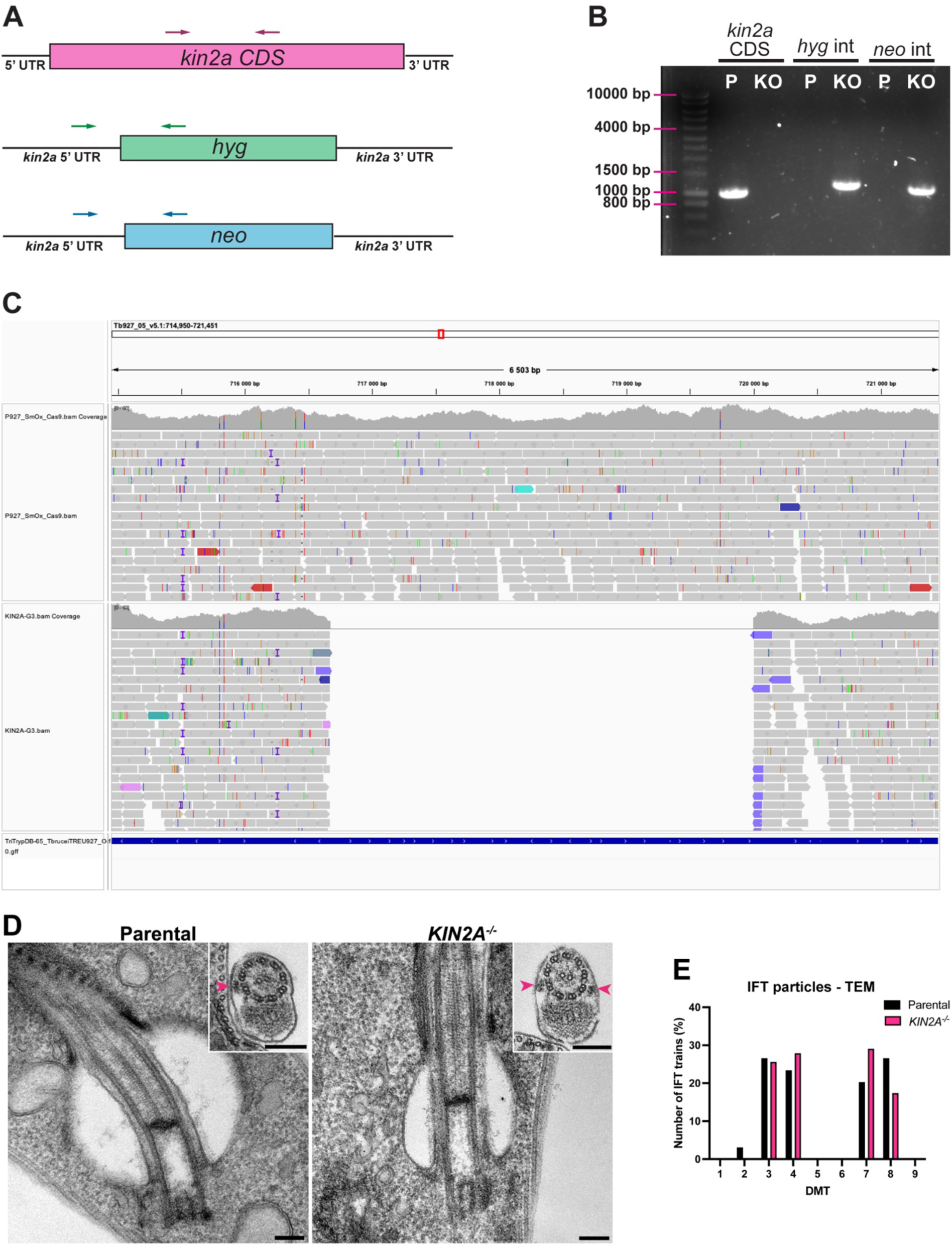
Validation of KIN2A deletion in *T. brucei*. **(A)** Schematic representation of the primers designed to verify the presence of the *kin2a* coding sequence (CDS) and the proper integration of hygromycin- or neomycin-resistance genes into the *kin2a* locus. **(B)** PCR products amplified from genomic DNA extracted from parental (+/+) and *KIN2A^-/-^* (-/-) cells, using the primers designed as shown in **A**. The gel confirms the absence of the *kin2a* CDS in *KIN2A^-/-^* cells and the correct integration of both resistance genes. **(C)** Whole-genome sequencing of parental and *KIN2A^-/-^* cells, demonstrating the complete deletion of the *KIN2A* gene. **(D)** Transmission electron microscopy (TEM) images of parental and *KIN2A^-/-^* cells. Transverse images of the flagellum base and cross-sections of the flagellum showing IFT trains (arrowheads) reveal no apparent structural differences between the two cell lines. Scale bars: 200 nm. **(E)** IFT trains positioned across the nine doublet microtubules (DMTs) of the axoneme from TEM images of parental and *KIN2A^-/-^* cells.

**Figure S6.**
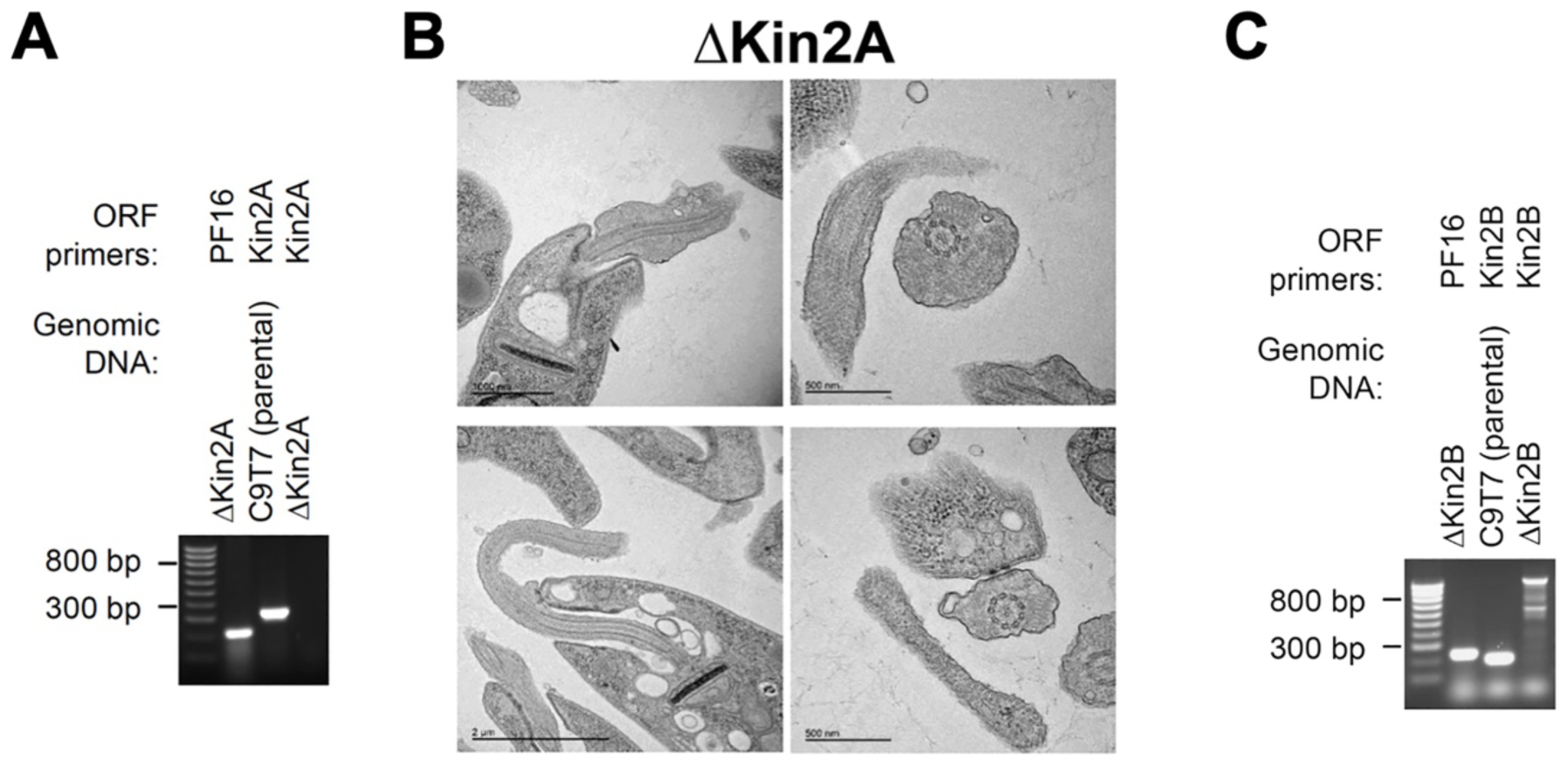
Deletion of KIN2A and KIN2B in *L. mexicana*. **(A, C)** PCR analysis of genomic DNA isolated from the indicated cell lines using primers for a control gene (*PF16*, a flagellar gene) and for KIN2A **(A)** or KIN2B **(C)**. (**B**) Transmission electron microscopy images showing that deletion of KIN2A does not visibly affect flagellar structure.

**Figure S7.**
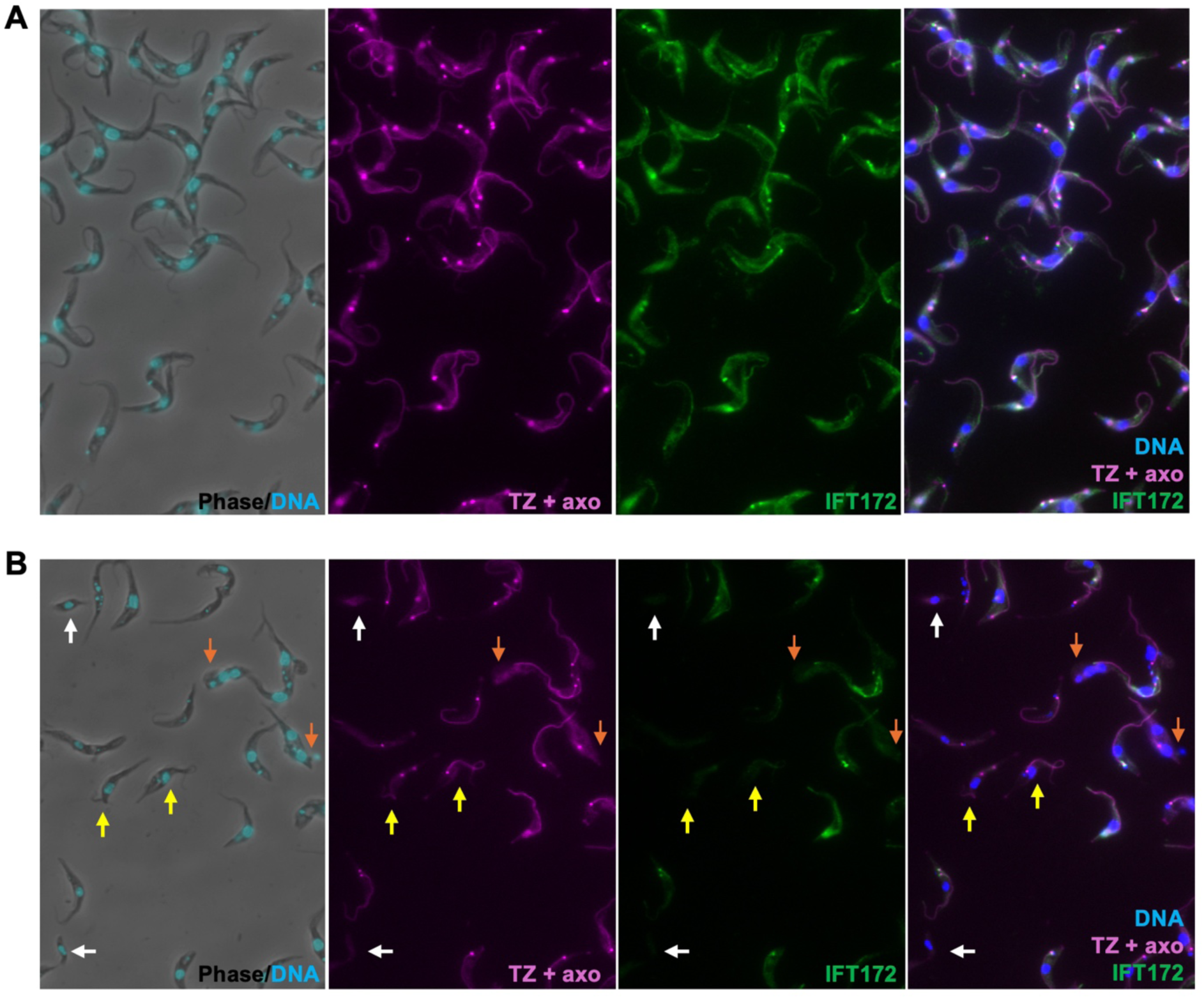
Validation of the Cas9-marker-free approach to disrupt the *IFT172* gene. Wild-type trypanosomes were transfected with a mixture containing Cas9, gRNA and repair templates introducing stop codons in each reading frame of the *IFT172* gene. Control cells **(A)** and Cas9-IFT172 **(B)** modified cells were grown for 48h, fixed and processed for IFA with mAb25 as marker of the axoneme (axo, magenta) and the anti-FTZC as marker of the transition zone (TZ, also in magenta), the antibody against IFT172 (green) and DAPI (blue). Examples of non-flagellated trypanosomes or cells with shorter flagella are indicated with the white and yellow arrows, respectively. Cells that duplicated their nucleus and kinetoplast but failed to grow a new flagellum are highlighted with orange arrows.

**Figure S8.**
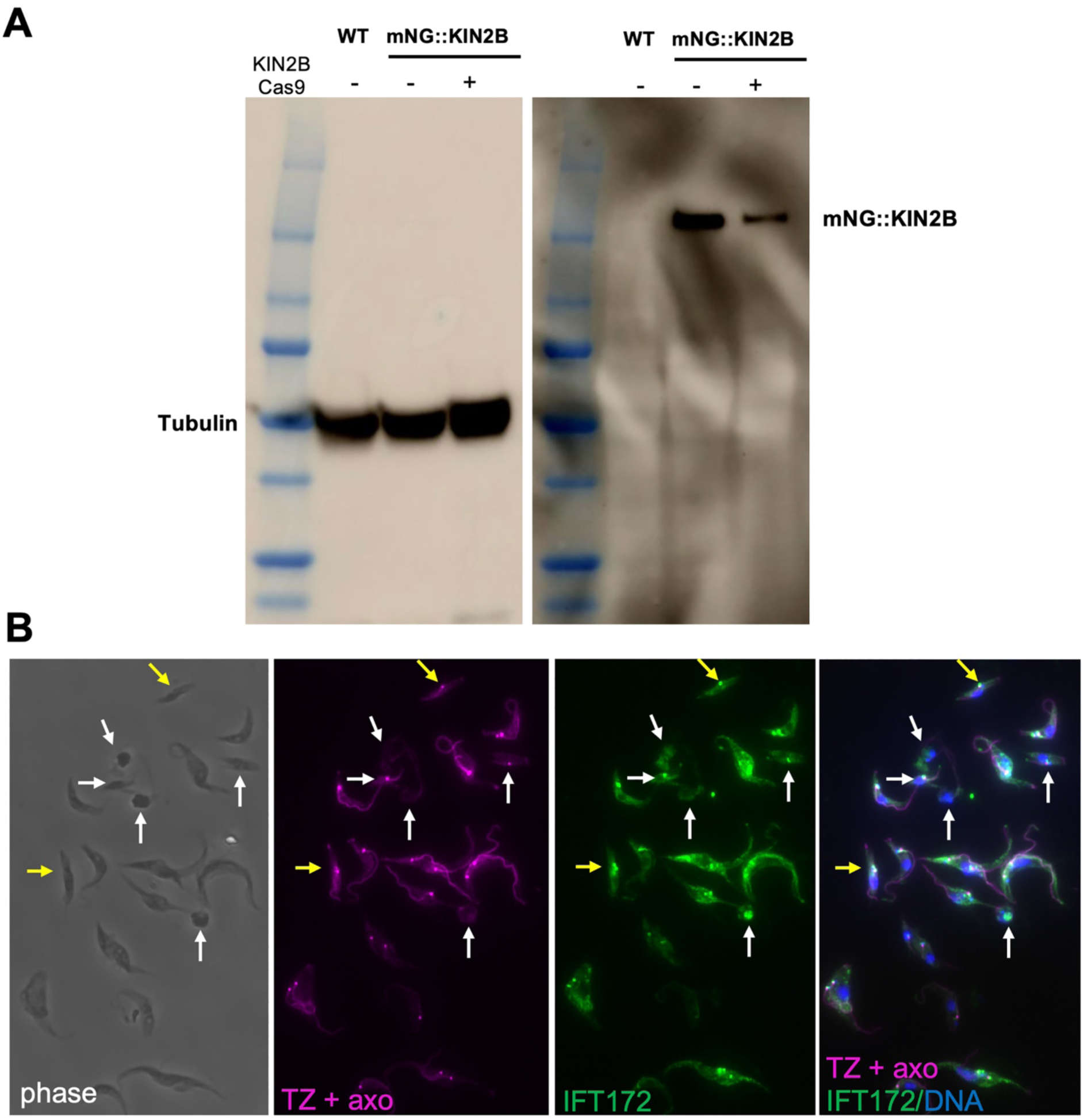
Inactivation of KIN2B inhibits flagellum formation and blocks IFT172 entry into the flagellum. Trypanosomes expressing mNG::KIN2B were nucleofected with a mixture containing Cas9, gRNA and repair templates introducing stop codons in each reading frame of the *KIN2B* gene. Cells were grown for 48h and samples were used for either **(A)** western blotting with the anti-Ty-1 tag BB2 antibody and the anti-tubulin TAT-1 as loading control or **(B)** fixed and processed for IFA with mAb25 as marker of the axoneme (axo, magenta) and the anti-FTZC as marker of the transition zone (TZ, also in magenta), the antibody against IFT172 (green) and DAPI (blue). Examples of non-flagellated trypanosomes or cells with shorter flagella are indicated with the white and yellow arrows, respectively.

**Video 1: Processive movement of the full-length and truncated KIN2A motors. (Top left panel)** The full-length KIN2A motor moves on surface-attached microtubules in two populations, a slower fraction and a faster fraction. The slower motors are highlighted by the yellow arrow and the faster ones by the white arrows. The full-length KIN2A^HALO^ motors are labelled using Alexa Fluor 488 dye. **(Top right panel)** The truncated KIN2A^1-690^ also exhibited a faster group of motors as pointed out by white arrows. Here the motors are labelled via a GFP-tag. **(Bottom left panel)** The artificially dimerized KIN2A^1-380^ GCN4 exhibits fast velocity as pointed out by white arrows. The KIN2A^1-380^ GCN4^HALO^ motors were labelled with Alexa Fluor 488 dye. The microtubules were labelled with ATTO550. Videos are sped up 1.47x. Scale bar: 5 µm.

**Video 2: Processive movement of the full-length and truncated KIN2B motors. (Left panel)** The full-length KIN2B motor moves on surface-attached microtubules as pointed out in white arrows. The full-length KIN2B^Halo^ motors are labelled using Alexa Fluor 660. **(Right panel)** The truncated KIN2B^1-378^ GCN4 exhibit faster movement compared to the full-length KIN2B as pointed out by white arrows. The KIN2B^1-378^ GCN4^HALO^ motors were labelled with Alexa Fluor 660. The microtubules were labelled with ATTO550. Videos are sped up 1.47x. Scale bar: 5 µm.

**Video 3:** The 30-second acquisition of the trypanosome shown in **Fig.5A** expressing IFT81::mNG. Scale bar: 5 µm.

**Video 4:** The 30-second acquisition of the trypanosome shown in **Fig.5C** expressing mNG::KIN2A. Scale bar: 5 µm.

**Video 5:** The 30-second acquisition of the trypanosome shown in **Fig.5E** expressing mNG::KIN2B. Scale bar: 5 µm.

**Video 6:** The 80-second acquisition of *L. mexicana* cells expressing either mNG::KIN2A (**left**) or mNG::KIN2B (**right**). The top panels show the raw fluorescence micrograph while the bottom ones show a background-subtracted image to enhance small structures.

**Video 7:** The 15-second acquisition of the cell shown in **Fig. 6A-B** expressing mNG::KIN2B and tdT::IFT140. The forked and triangular arrowheads indicate full-length particles and proximal KIN2B particles, respectively. Scale bar: 5 µm.

**Video 8:** The 30-second acquisition of the *KIN2A^-/-^* cell expressing mNG::IFT81 shown in **Fig. 8F**. Scale bar: 5 µm.

**Video 9:** The 30-second acquisition of the *KIN2A^-/-^* cell expressing mNG::KIN2B shown in **Fig. 8G**. Scale bar: 5 µm.

**Table S2:**
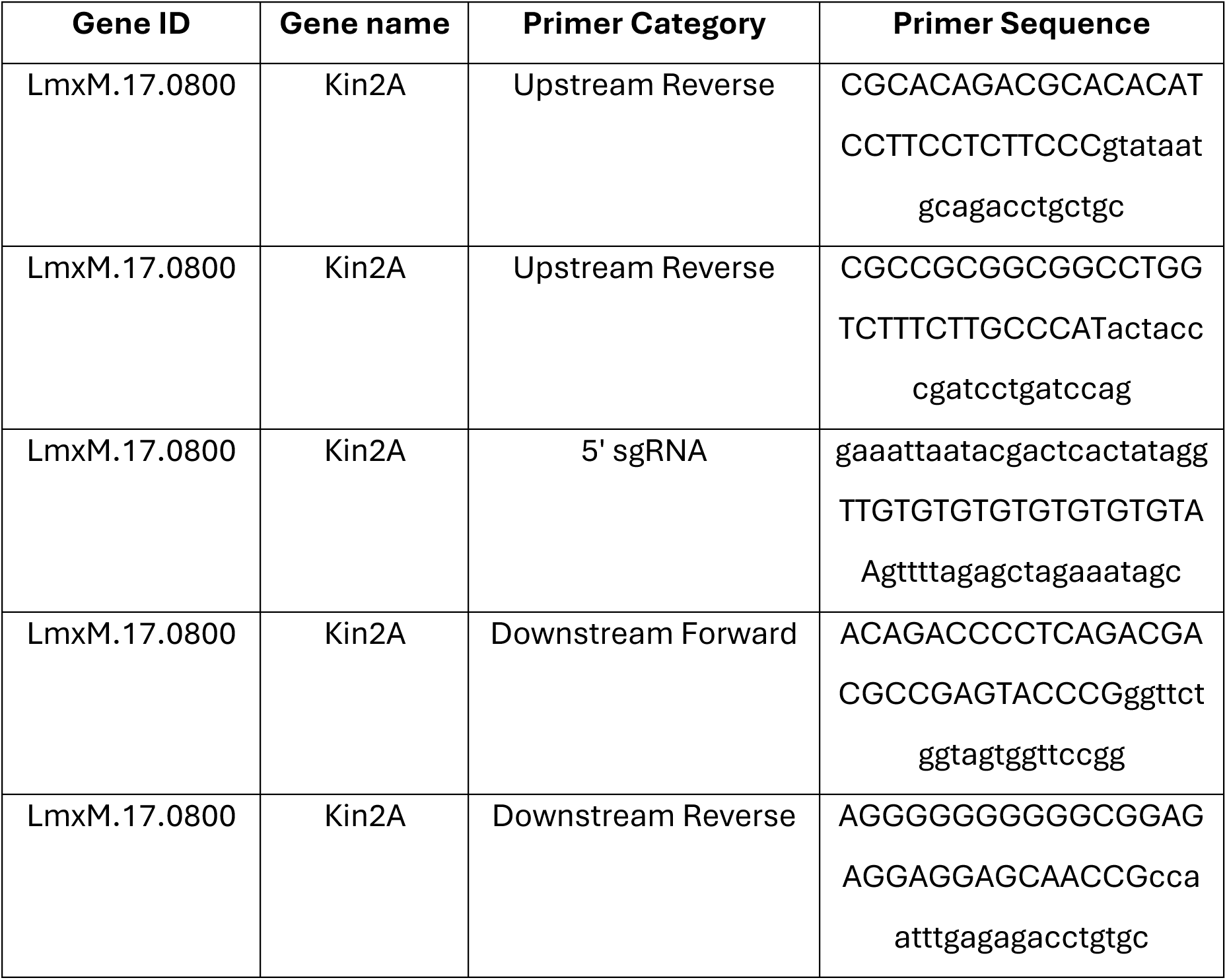
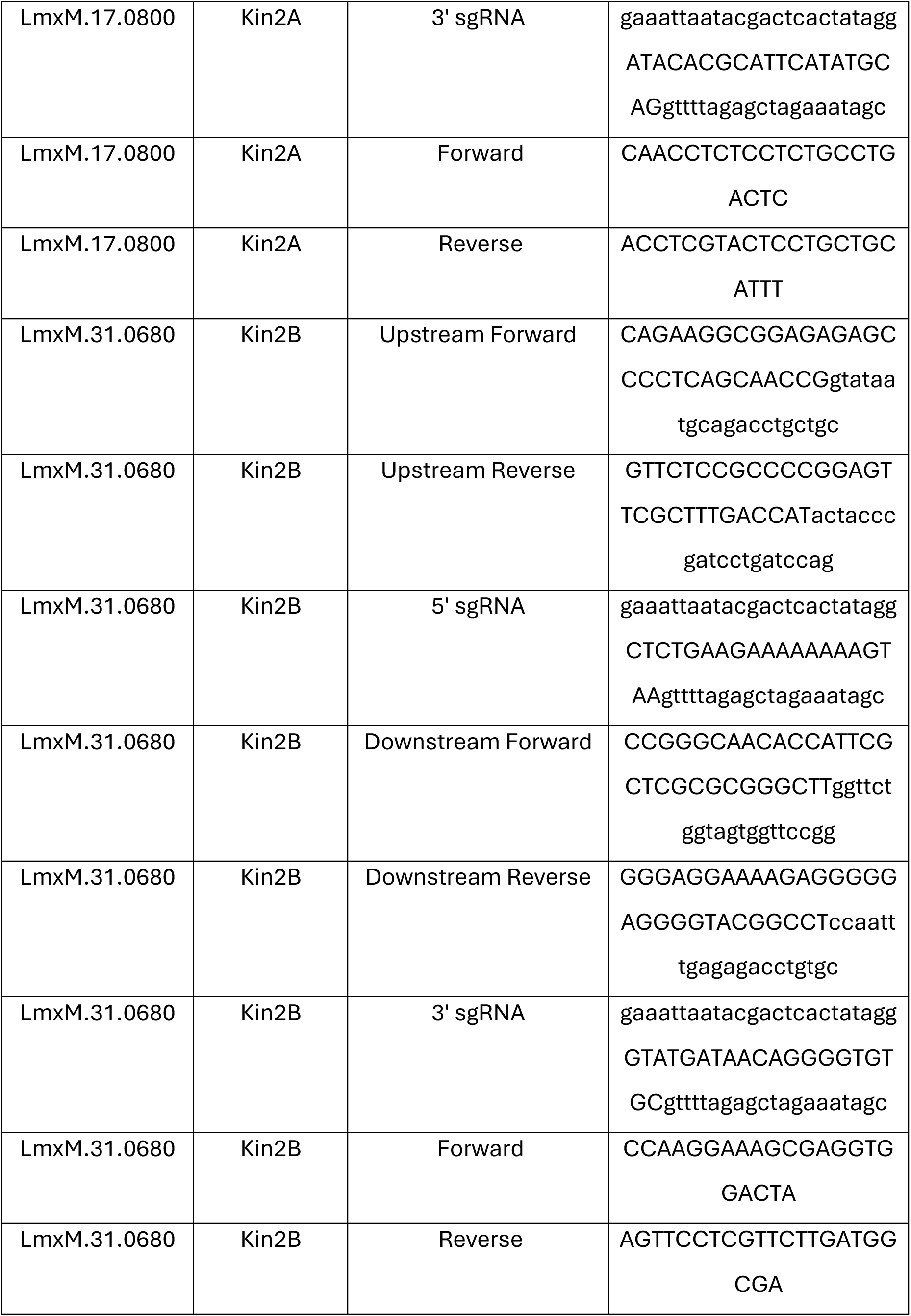
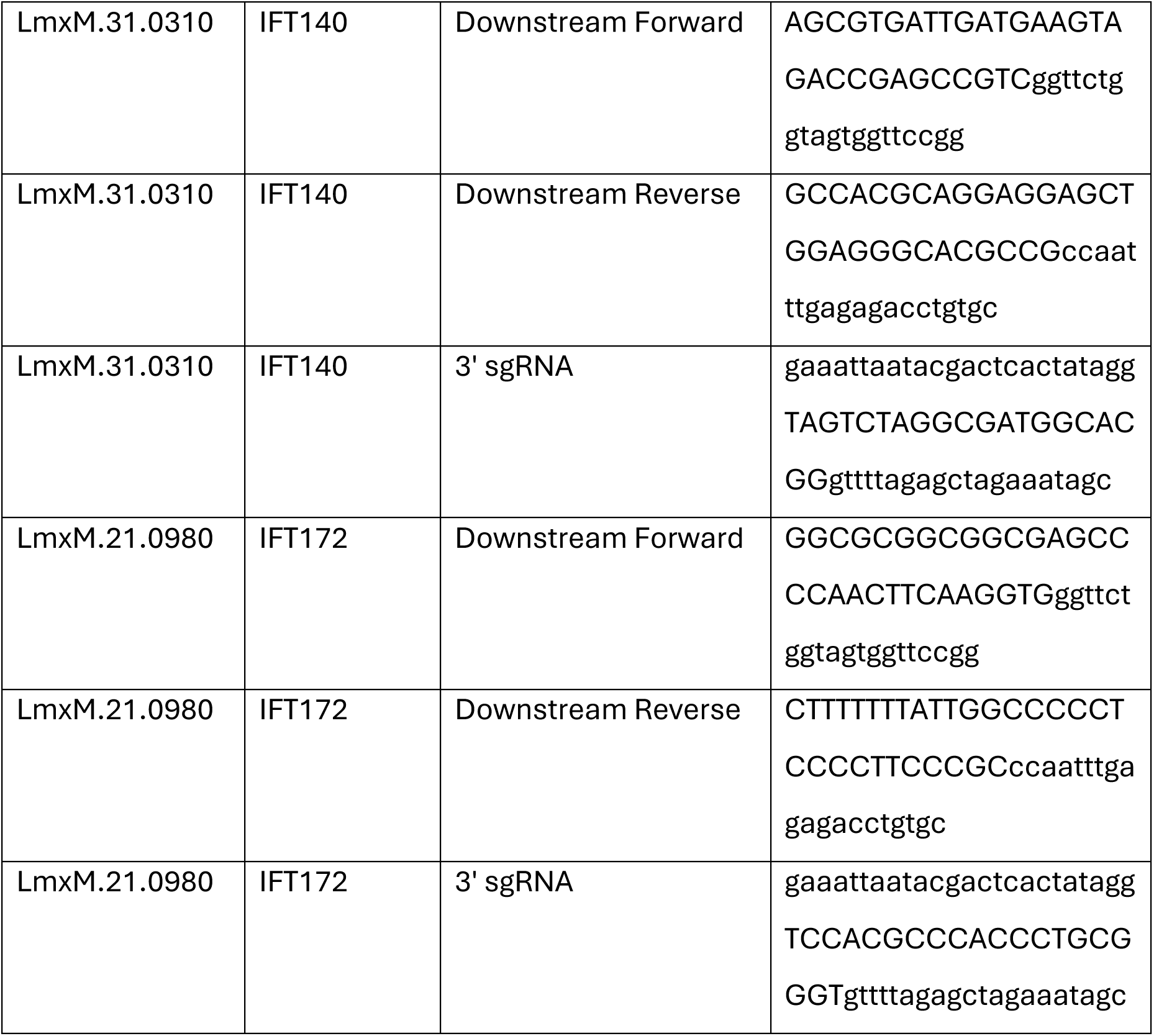
Sequence of the primers used in this study.

